# Dynamic epistasis analysis reveals how chromatin remodeling regulates transcriptional bursting

**DOI:** 10.1101/2021.12.15.472793

**Authors:** Ineke Brouwer, Emma Kerklingh, Fred van Leeuwen, Tineke Laura Lenstra

## Abstract

Transcriptional bursting has been linked to the stochastic positioning of nucleosomes. However, how bursting is regulated by remodeling of promoter nucleosomes is unknown. Here, we use single-molecule live-cell imaging of *GAL10* transcription in budding yeast to measure how transcriptional bursting changes upon single and double perturbations of chromatin remodeling factors, the transcription factor Gal4 and preinitiation complex (PIC) components. Using dynamic epistasis analysis, we reveal how remodeling of different nucleosomes regulates individual transcriptional bursting parameters. At the nucleosome covering the Gal4 binding sites, RSC acts synergistically with Gal4 binding to facilitate each burst. Conversely, nucleosome remodeling at the TATA box controls only the first burst upon galactose induction. In the absence of remodelers, nucleosomes at canonical TATA boxes are displaced by TBP binding to allow for transcription activation. Overall, our results reveal how promoter nucleosome remodeling, together with transcription factor and PIC binding regulates the kinetics of transcriptional bursting.

## Introduction

Transcription is a highly dynamic process that, for many genes, occurs in stochastic bursts of high transcriptional activity, interspersed by periods with no transcriptional activity. To achieve the correct transcriptional output of a bursting gene, transcription regulatory factors may modulate when a burst starts (burst frequency), when it ends (burst duration), and the rate of polymerase loading during a burst (initiation rate) (Bartman et al., 2016; Donovan et al., 2019a; Fukaya et al., 2016; Pimmett et al., 2021; Rodriguez et al., 2019; Tunnacliffe and Chubb, 2020; Zoller et al., 2018). Yet, it remains elusive how each of these steps is controlled. Previous studies from budding yeast have suggested a link between bursting and chromatin structure (Brown et al., 2013; Dadiani et al., 2013; Donovan et al., 2019a; Eck et al., 2020; Mehta et al., 2018; Shelansky and Boeger, 2020). Inference of transcriptional parameters from mRNA or protein distributions suggests that bursting is affected by mutations in different chromatin regulators (Mehta et al., 2018; Raser and O’Shea, 2004; Weinberger et al., 2012). In addition, single-cell mapping of nucleosome conformations at the *PHO5* promoter by methylation protection and by electron microscopy suggest stochastic transitions between different promoter configurations, of which only some may be permissive for transcription (Brown et al., 2013; Small et al., 2014). However, how different promoter nucleosome configurations affect the dynamics of transcription in living cells is unexplored.

The positioning of nucleosomes throughout the genome is controlled by chromatin remodeling enzymes. For promoter regions, the most important remodelers are RSC and SWI/SNF, which together maintain a promoter architecture consisting of a nucleosome-depleted region (NDR) flanked by two nucleosomes referred to as the +1 and -1 nucleosome (Kubik et al., 2019; Prajapati et al., 2020; Rawal et al., 2018). Upon depletion of RSC, the +1 nucleosome shifts into the NDR for 70% of all genes (Klein-Brill et al., 2019; Kubik et al., 2018). At highly expressed genes, SWI/SNF acts redundantly with RSC to maintain the NDR (Kubik et al., 2019; Rawal et al., 2018). In addition, RSC also regulates partially unwrapped unstable nucleosomes, referred to as fragile nucleosomes, that are often found in the promoters of highly expressed genes, such as *GAL10* (Brahma and Henikoff, 2019; Floer et al., 2010; Kubik et al., 2015; Weiner et al., 2010; Xi et al., 2011). The interplay between different remodelers at the same promoter is a highly dynamic process. Individual remodelers bind to chromatin with residence times of a few seconds and multiple remodelers can sequentially occupy the same promoter within minutes (Kim et al., 2021).

Nucleosomes affect different steps in the transcription activation process. For example, nucleosomes inhibit TF binding, and change the residence time of TFs that bind to nucleosomal DNA (Donovan et al., 2019b, 2019a; Luo et al., 2014; Mivelaz et al., 2020; Zhu et al., 2018). TF binding to nucleosomal DNA is facilitated by remodelers, as loss of RSC reduces TF occupancy, reduces TF binding frequency and increases TF residence time (Floer et al., 2010; Mehta et al., 2018). Besides influencing TF binding, movement of the +1 nucleosome in RSC depleted cells increases nucleosome coverage of the TATA box and transcription start site (TSS), which affects TBP binding, PIC assembly dynamics and transcription (Klein-Brill et al., 2019; Kubik et al., 2018; Nguyen et al., 2021; Wang et al., 2021). However, how each of these mechanisms influence transcriptional bursting is unclear.

In this study, we aim to dissect the role of promoter nucleosome remodeling in regulating the dynamics of transcriptional bursting. We acutely depleted the remodelers RSC and SWI/SNF and used single-molecule live-cell imaging to measure changes in transcription dynamics at the *GAL10* gene in budding yeast. To decipher the regulatory mechanisms of remodeling at specific promoter nucleosomes, we combined chromatin remodeler depletion with perturbations of the TF Gal4 and PIC components, and analyzed the effect of these single and double perturbations on each dynamic parameter of transcription using dynamic epistasis analysis. This analysis method identified synergistic effects between the different perturbations and revealed specialized functions of different nucleosomes on transcription dynamics. We found that the fragile nucleosome at the Gal4 binding sites is controlled by the redundant action of RSC and Gal4 binding to facilitate consecutive bursts of transcription. The nucleosomes around the TATA box and TSS are remodeled in a partially redundant manner by RSC and SWI/SNF, to allow for the first burst of transcription after activation. In addition, our results revealed a role for TBP in competing with nucleosomes at the TATA element to enable transcription in the absence of chromatin remodelers. Overall, our study exposed how remodeling at the different promoter nucleosomes controls the accessibility of the DNA for binding of transcription factors and the PIC, and how this impacts different kinetic parameters of transcriptional bursting.

## Results

### Remodeling of *GAL10* promoter nucleosomes at the UASs and the TSS by RSC affects induction time and time between bursts

Upon addition of the sugar galactose to yeast cells, nucleosomes in the promoter region of *GAL10* are remodeled by nucleosome remodeling complexes to allow for transcription activation (Bryant et al., 2008). MNase-seq with high and low MNase concentrations showed the coverage of both stable and fragile nucleosomes, respectively, in inactive and active conditions (Kubik et al., 2015) (Figure S1A-C). As reported previously (Bryant et al., 2008; Donovan et al., 2019a; Floer et al., 2010), in transcriptionally inactive conditions (raffinose), the *GAL10* promoter region showed three main nucleosomes: a fragile nucleosome at the Gal4 upstream activation sequences (UASs), and two well-positioned stable nucleosomes at the TATA box and the TSS (+1). Upon activation with galactose, the nucleosome covering the TATA box was evicted, and the +1 nucleosome was moved downstream, creating an NDR. Consistent with previous finding at wide NDRs, the fragile nucleosome at the UASs remained present(Brahma and Henikoff, 2019; Floer et al., 2010; Kubik et al., 2015).

To dissect how remodeling at each of these three nucleosomes regulates transcriptional bursting, we conditionally depleted chromatin remodelers from the nucleus, mapped the effect on the promoter nucleosome positioning, and linked this to their effect on transcription dynamics. First, we depleted the chromatin remodeling complex RSC, an important nucleosome remodeler controlling both the stable and fragile nucleosomes in promoter regions (Floer et al., 2010; Klein-Brill et al., 2019; Kubik et al., 2018). To map changes in nucleosome positions, we performed MNase-seq in galactose-rich media in cells where Sth1, the catalytic subunit of RSC, was depleted from the nucleus for 60 minutes using anchor-away (Haruki et al., 2008). Depletion of the essential protein Sth1 was confirmed by lack of growth on rapamycin-containing plates (Figure S2A) and by imaging of the mScarlet-anchor-away tagged Sth1 subunit (Figure S2B). In line with previous studies, acute RSC depletion lead to a fill-in and shortening of NDRs at the genome-wide level (Figure S1D,E) (Kubik et al., 2018; Rawal et al., 2018), as well as a slightly lower coverage of fragile nucleosomes in promoter regions (Figure S1,F,G). At the *GAL10* locus, RSC predominantly regulated fragile nucleosomes, with a lower coverage over the Gal4 UAS and the TSS (Figure 1B), in line with a previous report (Floer et al., 2010). In addition, a small shift in the stable +1 nucleosome around the TSS was observed (Figure 1A).

**Figure 1:**
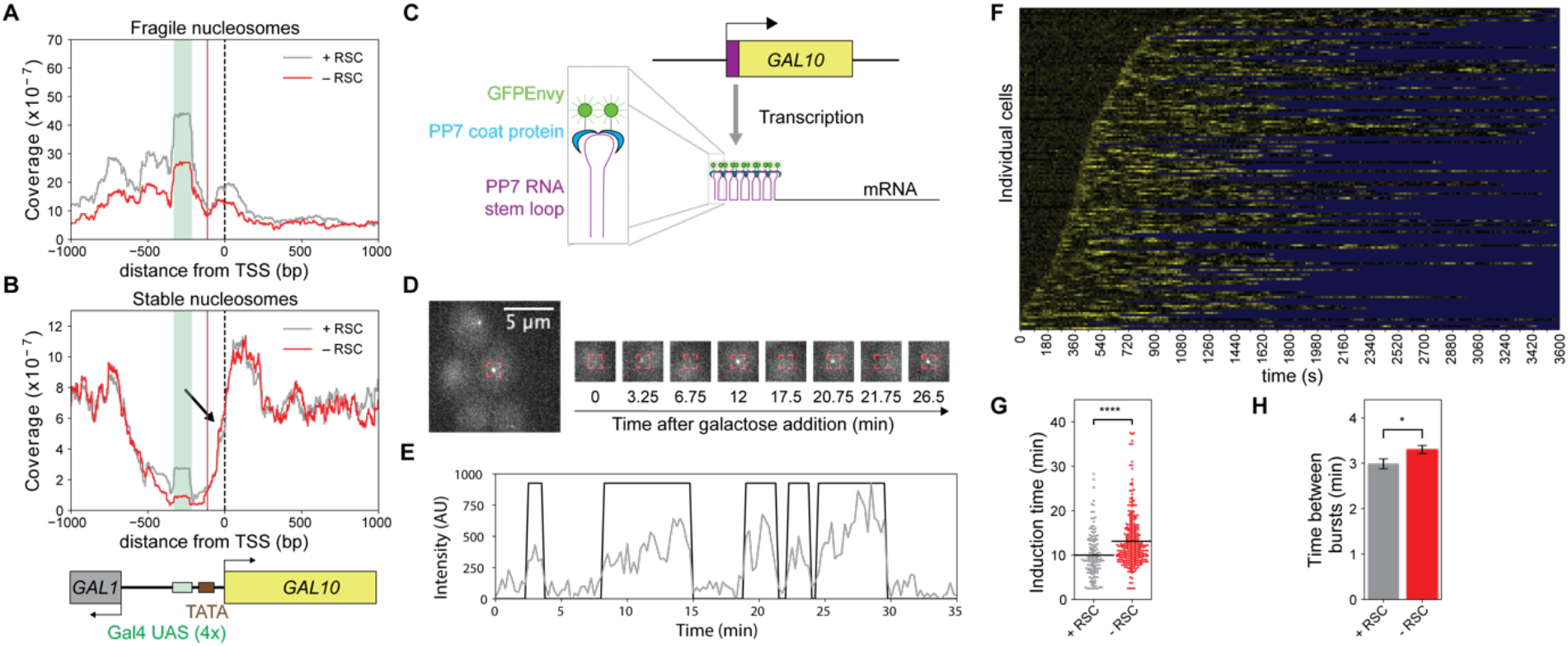
Remodeling of *GAL10* promoter nucleosomes at the UASs and the TSS by RSC affects induction time and time between bursts. (A) MNase-seq analysis of fragile nucleosomes in the *GAL10* promoter region showed a reduced coverage of fragile nucleosomes upon RSC depletion using anchor away of the catalytic subunit Sth1 by 60 min rapamycin treatment. (B) MNase-seq analysis of stable nucleosomes in the *GAL10* promoter region. Black arrow indicates a small shift of the +1 nucleosome into the nucleosome-depleted region (NDR) upon RSC depletion. MNase plots in (A) and (B) show one representative replicate out of two experiments. (C) Schematic of PP7 RNA labeling to visualize *GAL10* transcription in real-time. (D) Fluorescence signal of an individual transcription site (TS) in a representative cell over time. Red box: location of *GAL10* TS. (E) Quantification of TS intensity over time of the cell in (D) (grey) and binarization (black). (F) Heatmap of the TS intensity in N=143 cells (rows) in the presence of RSC. Yellow: fluorescence intensity of TS; blue: region excluded from analysis. (G) Cells that were depleted of RSC (red) showed an increased *GAL10* induction time compared to the non-depleted cells (grey). (H) Cells that were depleted of RSC (red) showed an increased time between burst of *GAL10* transcription compared to the non-depleted cells (grey). Error bars are standard deviation based on 1000 bootstrap repeats. Significance in (G) and (H) was determined by bootstrap hypothesis testing (MacKinnon, 2007); *: p < 0.05, ****: p < 0.00005. See also Figure S1, S2, S3, S4

To understand how this perturbed chromatin structure upon RSC depletion affects transcription dynamics of the *GAL10* gene, we employed the PP7 technology to directly monitor transcription in individual cells in real-time (Figure 1C, Video S1). *GAL10* was endogenously tagged with 14 PP7 repeats in the 5’UTR and upon transcription, the PP7 RNA stem loops are bound by PP7 coat protein fused to GFP-Envy (Larson et al., 2011). Using widefield fluorescence microscopy, the accumulation of RNAs at the transcription site (TS) was visualized as a bright spot in the nucleus (Figure 1D, Video S1), of which the intensity was tracked over time (Figure 1E,F). From these intensity traces, the parameters of transcriptional bursting were determined: the active fraction (the fraction of cells that show a TS within 1 hour after galactose addition), the induction time (the time between galactose addition and the first burst of transcription), the burst duration, the time between bursts (as a measure for burst frequency) and the burst intensity. The burst size, defined as the total number of RNAs produced during a burst, is dependent on the burst duration and the burst intensity. Upon RSC depletion, we observed an increase in induction time (Figure 1G) and in the time between bursts (Figure 1H), while the active fraction, burst duration, and burst intensity showed only minor or no changes (Figure S3A,H-J). Using single-molecule fluorescence in situ hybridization (smFISH), we validated that the effect of RSC depletion on steady-state transcription did not vary between the different cell-cycle stages (Figure S2C). Remodeling of the fragile *GAL10* promoter nucleosomes at the UASs and the TSS by RSC is thus important for regulating the induction time as well as the start of each burst of *GAL10* transcription.

### Remodeling of promoter nucleosome at the TATA box by SWI/SNF regulates *GAL10* induction time

Nucleosomes at promoters of highly expressed genes are remodeled by both RSC and SWI/SNF (Kubik et al., 2019; Rawal et al., 2018). Therefore, we subsequently addressed how SWI/SNF remodeling affects nucleosome positioning and transcription dynamics at *GAL10*, by nuclear depletion of Swi2, the catalytic subunit of SWI/SNF (Figure S2A,B). Consistent with previous reports, MNase-seq indicated that upon depletion of SWI/SNF the promoter nucleosome architecture did not change at the genome-wide level (Figure S1H,I). In contrast, for *GAL10*, the coverage within the NDR and specifically at the TATA element was increased (Figure 2A). Both genome-wide (Figure S1J,K) and at the *GAL10* locus (Figure S3K,L), no change in fragile nucleosomes was observed. At the level of transcription, depletion of SWI/SNF resulted in an increase in induction time (Figure 2D) but had a minor effect on the other *GAL10* transcriptional parameters (Figure 2G, S2C, S3B-E,H-J). Higher nucleosome coverage within the NDR and at the TATA box upon SWI/SNF depletion thus affects induction time, but once the cells are activated, this increased coverage has no effect on transcription.

**Figure 2:**
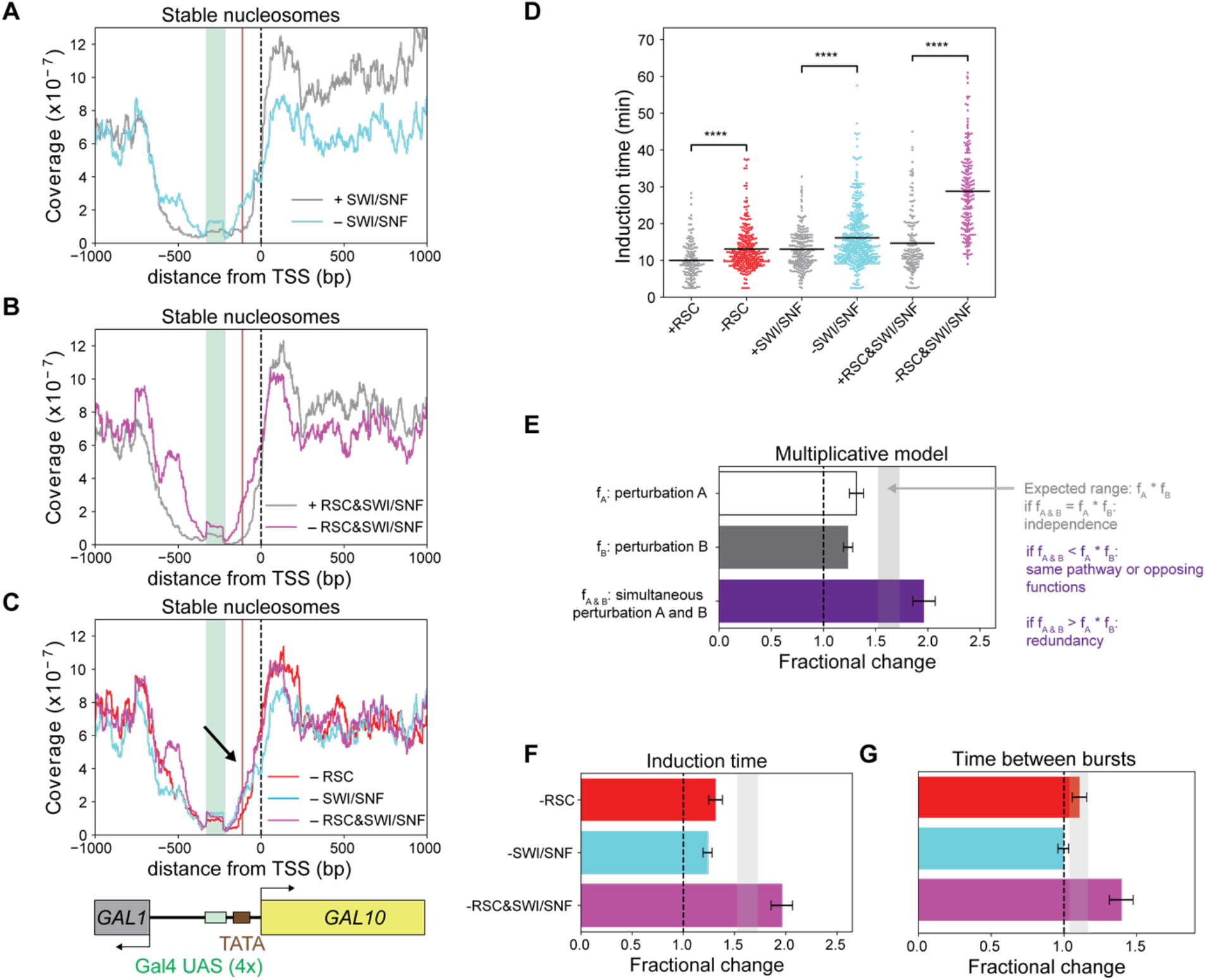
Partially redundant remodeling of nucleosomes by RSC and SWI/SNF at the TATA box and TSS synergistically affects induction time and time between bursts. (A) MNase-seq analysis in the *GAL10* promoter region showed higher coverage of stable nucleosomes around the TATA box upon depletion of SWI/SNF by anchor away of the catalytic subunit Swi2. (B) MNase-seq analysis of stable nucleosomes in the *GAL10* promoter region showed higher coverage at the TATA box and around the TSS upon simultaneous depletion of RSC and SWI/SNF than depletion of either RSC or SWI/SNF individually. (C) Overlay of MNase-seq analysis of stable nucleosomes in the *GAL10* promoter region upon depletion of RSC, SWI/SNF and simultaneous depletion of RSC and SWI/SNF showed increased coverage (black arrow). MNase plots in (A), (B) and (C) show one representative replicate out of two experiments. (D) Cells that were depleted of RSC and/or SWI/SNF (red, cyan, magenta) had an increased *GAL10* induction time compared to the non-depleted cells (grey). Significance determined by bootstrap hypothesis testing (MacKinnon, 2007); ****: p < 0.00005. (E) The multiplicative model for dynamic epistasis analysis was used to assess the effect of double perturbations. The expected effect of a double perturbation for independent processes is the product of the effect of the individual perturbations (grey bar). If the observed effect is smaller than this expected effect, the perturbations are in the same pathway or have opposing functions. If the observed effect is larger, the processes are redundant. (F) The increase in *GAL10* induction time when simultaneously depleting RSC and SWI/SNF was larger than expected based on their individual depletions. Grey bar: expected effect based on dynamic epistasis analysis. (G) The increase in time between *GAL10* transcriptional bursts in the RSC + SWI/SNF double depletion was larger than expected based on their individual depletions. Grey bar: expected effect based on dynamic epistasis analysis. Error bars in (F) and (G) are propagated standard deviations based on 1000 bootstrap repeats. See also Figure S1, S2, S3, S4

### Partially redundant remodeling of nucleosomes by RSC and SWI/SNF synergistically affects induction time and time between bursts

RSC and SWI/SNF have been shown to act redundantly, where in the absence of one, the other can take over its function (Kubik et al., 2019; Rawal et al., 2018). The effect of simultaneous depletion of both RSC and SWI/SNF on both chromatin structure and transcription dynamics is thus expected to be larger than the combined effect of their individual depletions. To test this, we simultaneously depleted both RSC and SWI/SNF using simultaneous anchor-away of Sth1 and Swi2 (Figure S2A,B). As expected, a larger effect on stable nucleosomes was observed than for either single depletion, both at the genome-wide level (Figure S1L,M) and the *GAL10* locus (Figure 2B,C), where the +1 nucleosome moved further into the NDR than either single depletion. However, the coverage at the TATA element of *GAL10* was similar to the coverage in the single SWI/SNF depletion, and the effect on fragile nucleosomes mimicked the effect in the single RSC depletion (Figure S1N,O,S3L). RSC and SWI/SNF are thus only partially redundant and have different functions to remodel promoter nucleosomes.

To interpret the effect of RSC&SWI/SNF double depletion in comparison to the effect of RSC or SWI/SNF single depletion, we performed epistasis analysis on each dynamic parameter of transcription and called this analysis method dynamic epistasis analysis. Dynamic epistasis analysis is based on classical epistasis analysis, a method to interpret phenotypic growth defects from genetic interactions, which allows one to determine whether mutant phenotypes act in the same or different pathways (van Leeuwen et al., 2017). Dynamic epistasis analysis compares the observed effect to the expected effect of a double perturbation on each dynamic parameter of transcription by taking the product of the fractional changes in this parameter observed in the individual perturbations. If two perturbations act independently, the double perturbation follows the expected effect (Figure 2E). Observing a larger effect than expected indicates that the two perturbed factors act on the same process and are at least partially redundant, while a smaller effect than expected indicates that the processes act in the same pathway or have opposing biochemical functions (van Leeuwen et al., 2017). For double depletion of RSC and SWI/SNF, a larger effect than expected was observed for both *GAL10* induction time and time between bursts (Figure 2D,F,G). In addition, the burst duration and intensity was shorter than expected, suggesting a smaller burst size in the double depletion (Figure S3H-J). As RSC and SWI/SNF are known to act redundantly (Kubik et al., 2019; Rawal et al., 2018), the observed synergistic effects for RSC and SWI/SNF validate our approach to identify functional epistatic relationships using transcription dynamics. In addition, these results showed that remodeling of the fragile nucleosomes at the UASs by RSC, and nucleosome displacement around the TATA element and the TSS by RSC and SWI/SNF, synergistically controls the induction time, time between bursts and burst size.

### Remodeling of promoter nucleosomes is important for multiple steps in gene activation

To understand in more detail how remodeling of promoter nucleosomes affects the kinetics of transcription activation, we estimated the number of rate-limiting steps from the induction time distributions by fitting a gamma distribution. The shape parameter *k* of such a fit is a measure of the number of rate-liming steps, under the assumption that the steps have equal rates. In the presence of remodelers, the distributions of induction times followed gamma distributions with *k* = ∼6 (Figure S4H,N,T), indicating that there are approximately 6 rate-liming steps during activation. However, single or double depletion of RSC and SWI/SNF before activation did not lead to a consistent change in *k* (Figure S4I,O,U), suggesting that the assumption of equal rates may no longer be valid. In addition, the high number of steps makes it challenging to extract a specific effect of remodeler depletion on the kinetics of induction. Likely, multiple rate-liming steps in the signaling pathway dominate and obscure any remodeler-specific steps of transcription activation. To expose remodeling-dependent activation steps, we attempted to eliminate signaling components by studying cells that are activated with galactose, then repressed with glucose-containing media, and subsequently reinduced with galactose. Consistent with previous reports, this re-induction is faster than the initial induction (Figure S4J,P,V), because cells have already adapted to galactose by cytoplasmic inheritance of the signaling proteins Gal3 and Gal1, referred to as transcriptional memory (Bryant et al., 2008; Kundu and Peterson, 2010). The observed re-induction times are now fit by a gamma distribution with *k* = ∼1 (Figure S4J,P,V) indicating that the re-induction process consists of a single rate-limiting step. Single depletion of either RSC or SWI/SNF before re-induction did not increase the number of steps (Figure S4K,Q), likely because of redundancy. However, simultaneous depletion of RSC and SWI/SNF increased the number of steps to ∼4 or ∼2, depending on the specific glucose-containing media used to repress before re-induction (Figure S4W,Y). This shift indicated that in the absence of remodelers, up to three remodeler-dependent activation steps are considerably prolonged and become rate-limiting (Figure S4Y). Remodeling by RSC and SWI/SNF thus regulates multiple steps in the activation pathway, perhaps by acting on the different promoter nucleosomes, each with their own effect on the kinetics.

Moreover, similar to the first induction, remodeler depletion during a second galactose exposure resulted in an increase in induction time and the time-between bursts. However, the lower burst duration and burst intensity observed in the double depletion during the first induction, was not reproduced during re-induction, suggesting that this effect may be context-specific (Figure S4E,G). For subsequent analysis, we therefore focused on the kinetics of induction and time between bursts. Overall, these memory experiments showed that remodeling of promoter nucleosomes by RSC and SWI/SNF promotes multiple activation steps.

### Transcription activation can occur in the absence of promoter nucleosome remodelers

The above experiments showed that simultaneous depletion of RSC and SWI/SNF reduces transcription, but a significant level of transcription was still observed (Figure 2D,F,G,B,S3H-J). Also, comparison of the nucleosome coverage in the double depletion (Figure 2B,C) to nucleosome positions in cells grown in raffinose, where *GAL10* is transcriptionally inactive (Figure S1A-C), indicated that even in the absence of both RSC and SWI/SNF, the nucleosome configuration changed upon galactose induction. Under these conditions, nucleosomes must thus be remodeled by a mechanism independent of RSC and SWI/SNF. This nucleosome remodeling could either be performed by other remodelers, or potentially by the transcriptional machinery itself. Although at the genome-wide level promoter nucleosomes are regulated mainly by RSC and SWI/SNF (Kubik et al., 2019), it is possible that specifically at *GAL10* one of the other remodeling complexes plays a role. To test this hypothesis, we performed smFISH to detect changes in *GAL10* nascent transcription upon individual depletion of the catalytic subunit of each of the seven remodeling complexes present in budding yeast with anchor-away (Figure S3P,Q). Apart from RSC, none of the remodeling complexes had a large effect on steady-state transcription levels of *GAL10*. We conclude that if other remodelers regulate nucleosomes in the *GAL10* promoter, they must act redundantly with RSC and SWI/SNF. Alternatively, the transcriptional machinery itself may have a potential role in perturbing nucleosome architecture.

### Remodeling of the fragile nucleosome at the UASs by RSC and Gal4 binding synergistically regulates time between bursts

The remodeler depletion experiments above allowed us to link nucleosome changes to changes in transcription dynamics, but both RSC and SWI/SNF depletion affected multiple nucleosomes simultaneously. To decipher the mechanism by which remodeling at individual nucleosomes controls transcriptional bursting, we combined remodeler depletions with perturbations that affect binding of regulatory factors at specific nucleosome positions. First, we focused on the fragile nucleosome covering the Gal4 UASs. Because the MNase-seq profiles in the *GAL10* promoter (Figure 1A,B, 2A-C and S3K,L) showed a change in occupancy of the fragile nucleosomes at the Gal4 UASs upon RSC depletion, but no effect upon SWI/SNF depletion, the fragile nucleosome at the UASs appears to be solely remodeled by RSC. The effect of remodeling of this nucleosome on the binding of Gal4 to the UASs and transcription dynamics were assessed in *GAL4/gal4Δ* cells, where the concentration of Gal4 was reduced, which is expected to lower the on-rate of Gal4 binding to the UASs (Figure 3A). In these cells with reduced Gal4 levels, an increase in induction time was observed (Figure 3B,C) with a modest effect on other transcriptional parameters (Figure 3D, S3F,G,M-O). Upon additional depletion of RSC (Sth1) in these *GAL4/gal4Δ* cells, the induction time was within the expected range, suggesting independent roles of Gal4 and RSC during the first Gal4 binding event (Figure 3C). In contrast, we observed a synergistic increase in the time between bursts (Figure 3D), which indicates that remodeling of the fragile nucleosome at the Gal4 UASs by RSC is redundant with binding of Gal4. Gal4 binding can thus substitute for RSC, possibly by partially unwrapping the nucleosome at the Gal4 UASs. Furthermore, this combined action of RSC remodeling and Gal4 binding at the fragile nucleosome needs to occur before the start of each burst of transcription.

**Figure 3:**
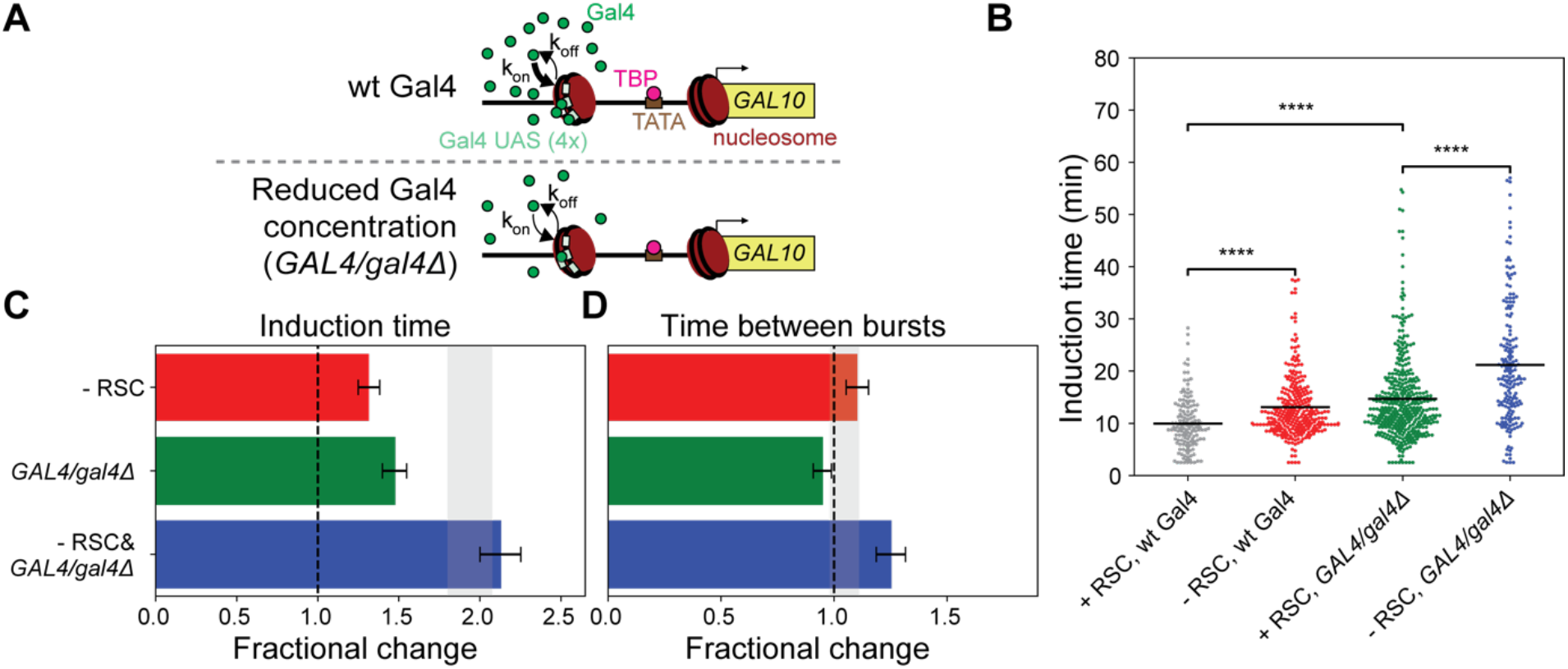
Remodeling of the fragile nucleosome at the UASs by RSC and Gal4 binding synergistically regulates time between bursts. (A) Schematic showing reduced Gal4 on rate in a *GAL4/gal4Δ* strain where Gal4 protein concentration was reduced. (B) *GAL10* induction time was increased in *GAL4/gal4Δ* cells and in RSC-depleted cells. Significance determined by bootstrap hypothesis testing (MacKinnon, 2007); ****: p < 0.00005. (C) Increase in *GAL10* induction time in RSC-depleted *GAL4/gal4Δ* cells was as expected based on individual perturbations. Grey bar: expected effect based on dynamic epistasis analysis. (D) The time between consecutive bursts of *GAL10* transcription in *GAL4/gal4Δ* cells increased more than expected based on individual perturbations. Grey bar: expected effect based on dynamic epistasis analysis. Error bars in (C) and (D) are propagated standard deviations based on 1000 bootstrap repeats. See also Figure S2, S3

### Nucleosome displacement at the TATA box and TSS by remodelers and TBP binding synergistically regulates induction time

Next, we focused on the nucleosomes in the *GAL10* promoter around the TATA and the TSS, a region crucial for PIC assembly. The first step in PIC assembly is TBP binding to the TATA box, which subsequently recruits the rest of the transcription machinery around the TSS to start transcribing the gene. The TATA box is covered by a nucleosome in inactive conditions, and gets exposed upon activation by galactose signaling (Figure S1A-C). Eviction of this nucleosome was reduced in SWI/SNF depleted cells. In addition, movement of the +1 nucleosome into the NDR after RSC depletion has been shown to affect TBP binding to the TATA box (Kubik et al., 2018). To uncover the mechanisms by which chromatin remodeling of these nucleosomes affect TBP binding and regulate transcription dynamics, we combined remodeler depletion with partial TBP depletion by introducing the anchor-away tag on one of the two alleles of the gene encoding for TBP in diploid cells, reducing the on-rate of TBP binding to the TATA element (Figure 4A). This partial depletion approach was chosen over full depletion because full depletion of TBP lead to a near-complete loss of *GAL10* transcription (Figure S5G). Although imaging indicated considerable TBP depletion (Figure S2B), we noted that even full TBP depletion was likely not complete, as evidenced by growth on rapamycin-containing plates of a TBP-FRB haploid strain (Figure S2A). Partial TBP depletion of one of two copies, however, only had a modest effect on *GAL10* transcription (Figure 4B,C,D,S5A,B,G-J), likely because the strong TATA box at *GAL10* ensured sufficient TBP binding even at a reduced TBP concentration. To study how nucleosome remodeling at the TATA box by RSC affects TBP binding, TBP and RSC were depleted simultaneously, by anchor-away of TBP and Sth1. This combined depletion resulted in a synergistic delay of *GAL10* induction (Figure 4B,C) and the expected effect on time between bursts (Figure 4D). A similar synergistic delay of indication time was obtained for SWI/SNF depletion in combination with partial TBP depletion (Figure S5E,F,K-P). In contrast to the fragile nucleosome at the Gal4 UASs, which requires remodeling before each burst of transcription, remodeling of the nucleosomes around the TATA box is mostly required to allow initial TBP binding and initiating the first burst of *GAL10* transcription and is not required for initiating subsequent bursts.

**Figure 4:**
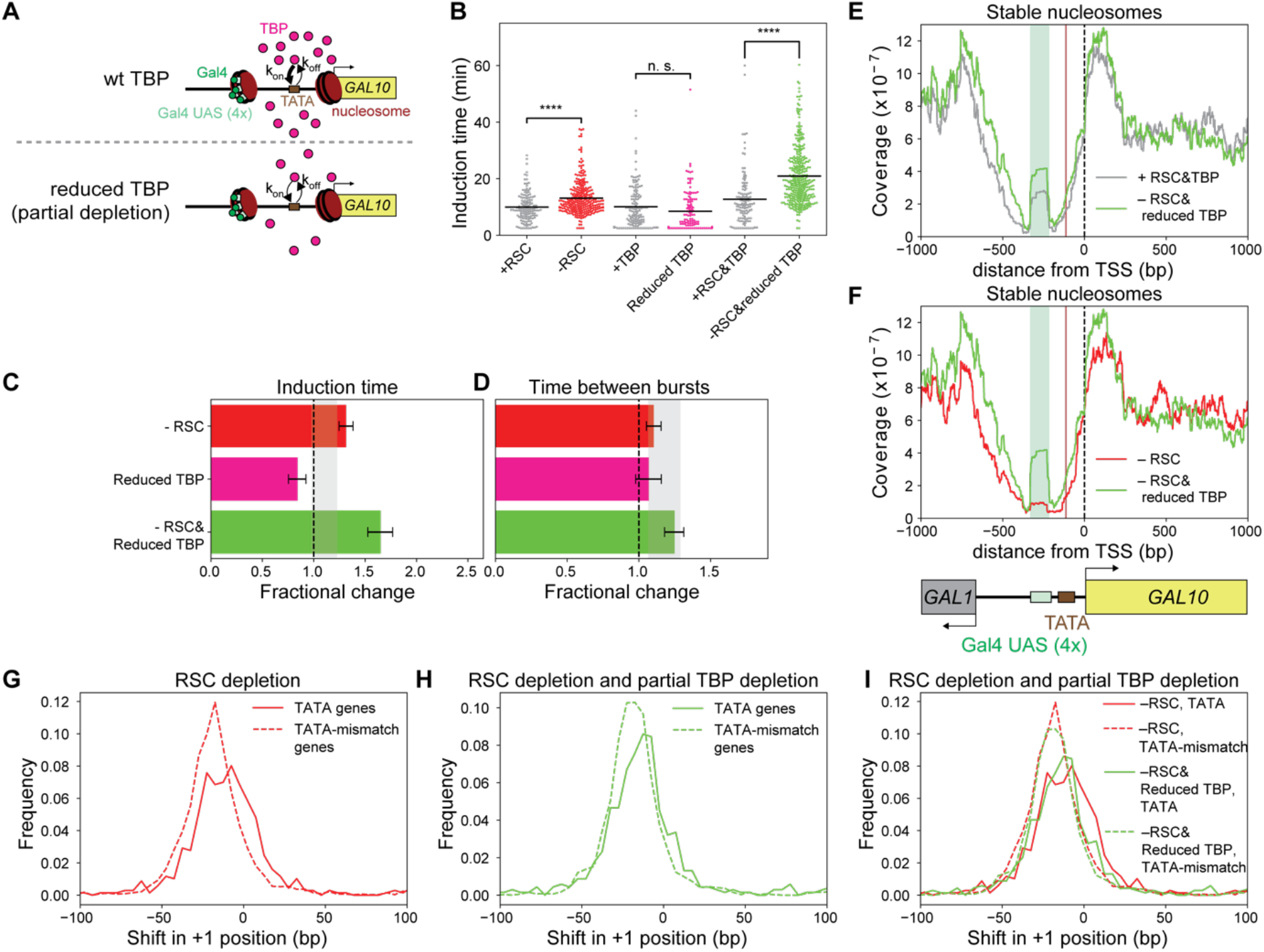
The nucleosomes at the TATA element and TSS are redundantly displaced by RSC and TBP to regulate the first burst of *GAL10* transcription. (A) Schematic showing reduced TBP on rate after addition of rapamycin in a diploid yeast strain where one of the two copies of TBP was depleted by anchor away. (B) *GAL10* induction time increased upon RSC depletion, did not change upon partial TBP depletion, and increased more in simultaneous RSC and partial TBP depletion. Significance determined by bootstrap hypothesis testing (MacKinnon, 2007); n.s.: not significant; ****: p < 0.00005. (C) The increase in *GAL10* induction time when simultaneously depleting RSC and partial TBP was larger than expected based on their individual depletions. Grey bar: expected effect based on dynamic epistasis analysis. (D) The time between consecutive bursts of *GAL10* transcription increased as expected in the double depletion of RSC and partial TBP. Grey bar: expected effect based on dynamic epistasis analysis. Error bars in (C) and (D) are propagated standard deviations based on 1000 bootstrap repeats. (E) MNase-seq analysis of stable nucleosomes in the *GAL10* promoter region showed increased coverage around the TSS and TATA when RSC and TBP were depleted. (F) MNase-seq analysis of stable nucleosomes in the *GAL10* promoter region showed higher coverage around the TATA and TSS when depleting RSC and TBP than when depleting only RSC. (G) Histogram of shift in +1 nucleosome upon depletion of RSC for both TATA genes and TATA-mismatch genes, showing that TATA-mismatch genes showed a larger shift in the +1 nucleosome than TATA genes. (H) Histogram of shift in +1 nucleosome upon simultaneous depletion of RSC and TBP for both TATA genes and TATA-mismatch genes. (I) Overlay of histograms of shift in +1 nucleosome upon depletion of RSC or simultaneous depletion of RSC and TBP for both TATA genes and TATA-mismatch genes, showing that partial TBP depletion resulted in a larger shift specifically at TATA-containing genes. Plots in (E-I) show one representative replicate out of two experiments. See also Figure S2, S5, S6

### In the absence of chromatin remodeling complexes, nucleosomes at the TATA box and TSS are evicted by TBP binding

The observed synergy between reduced nucleosome remodeling at the TATA box and lower TBP on-rate suggested a redundant role for TBP in affecting nucleosome positions. To test this hypothesis, MNase-seq was performed in cells depleted of RSC and partially depleted of TBP. Indeed, a larger shift in the +1 nucleosome and more nucleosome density over the TATA element in the *GAL10* promoter were observed in these double depleted cells than in RSC-depleted cells with wildtype TBP levels (Figure 4E,F,S6A-F). Thus, in cells with impaired chromatin remodeling, TBP binding at the TATA element contributes to positioning of the +1 nucleosome and the nucleosome covering the TATA box.

A possible function for TBP in nucleosome positioning was further supported by genome-wide analysis of our MNase-seq data of RSC-depleted cells. For genes with a canonical TATA motif, we observed a smaller shift in +1 nucleosome upon RSC depletion and less coverage at the TATA element than for genes lacking the canonical TATA motif (Figure 4G,S6G). Upon additional partial depletion of TBP, the ability of TBP to compete with nucleosomes is specifically reduced at genes with a strong TATA box, but not at TATA-mismatch genes (Figure 4H-I,S6H-I). These findings suggests that a strong TBP-TATA interaction is required for TBP to compete with nucleosomes.

### The ability of TBP to compete with nucleosomes requires a long residence time for TBP at the TATA element

To test whether the strength of the TBP-TATA interaction determines the ability of TBP to compete with nucleosomes, we perturbed the TBP-TATA interaction by mutating the TATA box in the *GAL10* promoter (Figure 5A) (Blake et al., 2006). Mutation of the TATA box reduced the burst duration and intensity (Figure S7A,B,K,L), which corroborated predictions of reduced burst size from fitting protein distributions with a theoretical bursting model to infer transcriptional parameters (Blake et al., 2006). In addition, the TATA box mutation increased the induction time and time between bursts (Figure 5B,D,E). Remarkably, depletion of RSC (Sth1) in these TATA-mutant cells resulted in two distinct cell populations, as evidenced by a large reduction in the fraction of cells that activated transcription during our 1-hour imaging experiment (Figure 5C). One population was not able to activate *GAL10* transcription at all likely because a nucleosome covering the TATA box could no longer be remodeled by TBP in the absence of RSC. For the other population, there was no additional effect of RSC depletion in terms of induction time (Figure 5D) and the time between bursts showed the expected effect (Figure 5E). This active population likely had no nucleosome covering TATA element that required remodeling, allowing TBP to bind and initiate transcription, resulting in RSC-independent induction of *GAL10* transcription. These results thus confirm our prediction that a strong TATA box is required for TBP to compete with nucleosomes.

**Figure 5:**
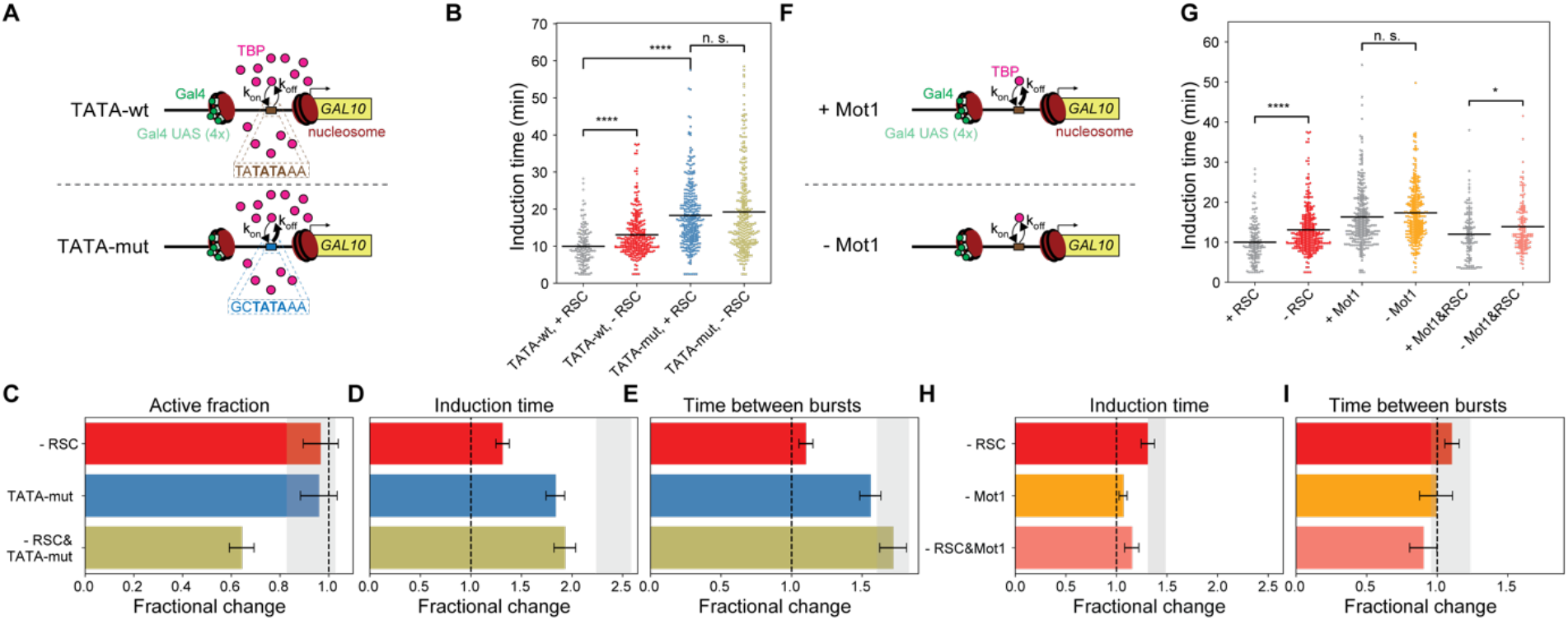
Competition between the nucleosome at the TATA and TBP depends on TBP residence time. (A) Schematic showing the reduced residence time of TBP upon mutating the TATA element in the *GAL10* promoter. (B) *GAL10* induction time was increased upon RSC depletion in cells with wt TATA element, but not in cells with mutated TATA element. Significance determined by bootstrap hypothesis testing (MacKinnon, 2007); n.s.: not significant; ****: p < 0.00005. (C) Upon RSC depletion in TATA-mutant cells, a fraction of cells was no longer able to activate *GAL10* transcription. Grey bar: expected effect based on dynamic epistasis analysis. (D) Mutating the TATA element increased *GAL10* induction time, but upon RSC depletion, there was no additional effect on the induction time. The effect of depletion of RSC in TATA-mutant cells was less than expected based on individual perturbations. Grey bar: expected effect based on dynamic epistasis analysis. (E) The effect on the time between consecutive bursts of *GAL10* transcription was as expected based on individual perturbations. Grey bar: expected effect based on dynamic epistasis analysis. (F) Schematic showing increased TBP residence time at the TATA element upon Mot1 depletion. (J) Mot1 depletion did not affect *GAL10* induction time. Upon simultaneous RSC and Mot1 depletion, there was a small effect on *GAL10* induction time. Significance determined by bootstrap hypothesis testing (MacKinnon, 2007); n.s.: not significant; *: p < 0.05; ****: p < 0.00005. (K) The effect of depletion of simultaneous RSC and Mot1 depletion was less than expected based on individual perturbations, where Mot1 rescued the effect of RSC depletion. Grey bar: expected effect based on dynamic epistasis analysis. (L) The effect on the time between consecutive bursts of *GAL10* transcription was as expected based on individual perturbations. Grey bar: expected effect based on dynamic epistasis analysis. Error bars in (C), (D), (E), (H) and (I) are propagated standard deviations based on 1000 bootstrap repeats. See also Figure S2, S7

The inability of TBP to compete with nucleosomes upon a TATA mutation indicated that TBP requires a longer residence time on DNA to compete with the nucleosome and activate transcription. We therefore reasoned that increasing the residence time of TBP would have the opposite effect to the TATA mutation, and should allow for increased ability of TBP to compete with nucleosomes. We tested this hypothesis by increasing the stability of TBP interaction with the TATA element by depletion of Mot1, the protein that facilitates TBP removal (Andrau et al., 2002; Dasgupta et al., 2002; Zentner and Henikoff, 2013) (Figure 5F). Single depletion of Mot1 had only a modest transcriptional phenotype (Figure 5G-I,S2K,S7C-F,M-O), perhaps because depletion was substantial but not complete, as indicated by growth of the Mot1 anchor-away strain on rapamycin-containing plates and imaging (Figure S2A,B). In agreement with our prediction, simultaneous depletion of RSC and Mot1 by anchor-away of Sth1 and Mot1, led to a rescue of the effect of RSC depletion on *GAL10* induction time and time between bursts. The stabilized TBP-TATA element interaction thus enhances the nucleosome competition ability of TBP, and is sufficient to activate transcription efficiently, even if in the absence of RSC remodeling.

### Nucleosome remodeling of RSC at the TSS acts antagonistically with Taf1 binding to control induction time

To test if the redundant nucleosome displacement at the TATA box by RSC and TBP binding is specific for the interaction between TBP and the TATA element rather than a general effect from potentially impaired PIC assembly by perturbed TBP-TATA interactions, we measured the effect of Taf1 depletion in combination with RSC depletion. Taf1 is part of the PIC but is recruited after TBP binding and has no direct interaction with the TATA element or the nucleosome at the TATA element, but rather interacts with the +1 nucleosome around the TSS (Joo et al., 2017) (Figure 6A). For Taf1-depleted cells, a delayed *GAL10* induction was observed (Figure 6B-D, Figure S7G-J,P-R). Rather than the synergistic effect observed between RSC and TBP depletion, we observed a smaller effect on induction time than expected when depleting RSC (Sth1) and Taf1, suggesting opposing roles of RSC and Taf1 in controlling induction time (Figure 6C). Therefore, in contrast to TBP, Taf1 cannot compete with nucleosomes. Conversely, nucleosome remodeling of RSC at the TSS acts antagonistically with Taf1 binding to control induction time.

**Figure 6:**
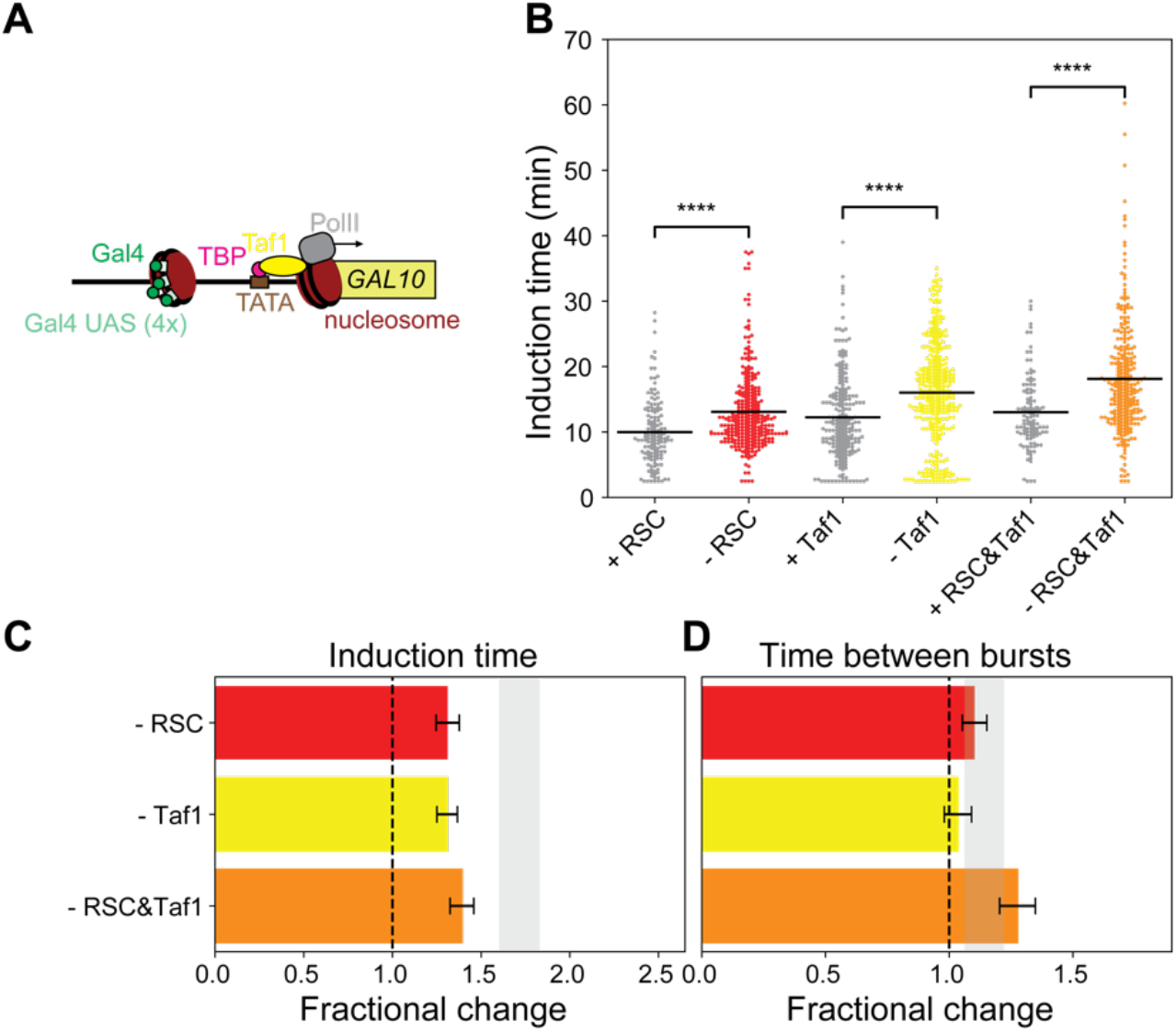
Antagonistic effect on induction time by RSC and Taf1 shows that nucleosomes cannot be competed away by Taf1. (A) Schematic showing Taf1 binding at the +1 nucleosome in the *GAL10* promoter region. (B) Taf1 depletion and simultaneous RSC and Taf1 depletion increased induction time of *GAL10*. Significance determined by bootstrap hypothesis testing (MacKinnon, 2007); ****: p < 0.00005. (C) Taf1 and RSC depletion had a smaller effect on induction time than expected based on the individual depletions, indicating that RSC and Taf1 have opposing functions. Grey bar: expected effect based on dynamic epistasis analysis. (D) Taf1 and RSC depletion had the expected effect on time between consecutive bursts. Grey bar: expected effect based on dynamic epistasis analysis. Error bars in (C) and (D) are propagated standard deviations based on 1000 bootstrap repeats. See also Figure S2

## Discussion

In this study, we use a dynamic epistasis analysis of single and double perturbations of nucleosome remodeling and the transcription machinery in combination with nucleosome mapping experiments and single-molecule live-cell imaging at the *GAL10* gene in *S. cerevisiae, to* uncover how remodeling of promoter nucleosomes regulates transcriptional bursting. Based on our findings, we propose a model (Figure 7) where different promoter nucleosomes have specialized roles in controlling transcriptional dynamics. Specifically, a fragile nucleosome covering the Gal4 UASs is repeatedly remodeled by RSC, in a manner redundant with Gal4 binding, which regulates the correct start of each transcriptional burst. Additionally, nucleosomes around the TATA element and the TSS are positioned by an interplay of RSC, SWI/SNF and TBP to allow for transcription of *GAL10*. This latter activity is only rate-limiting in the first burst of *GAL10* transcription.

**Figure 7:**
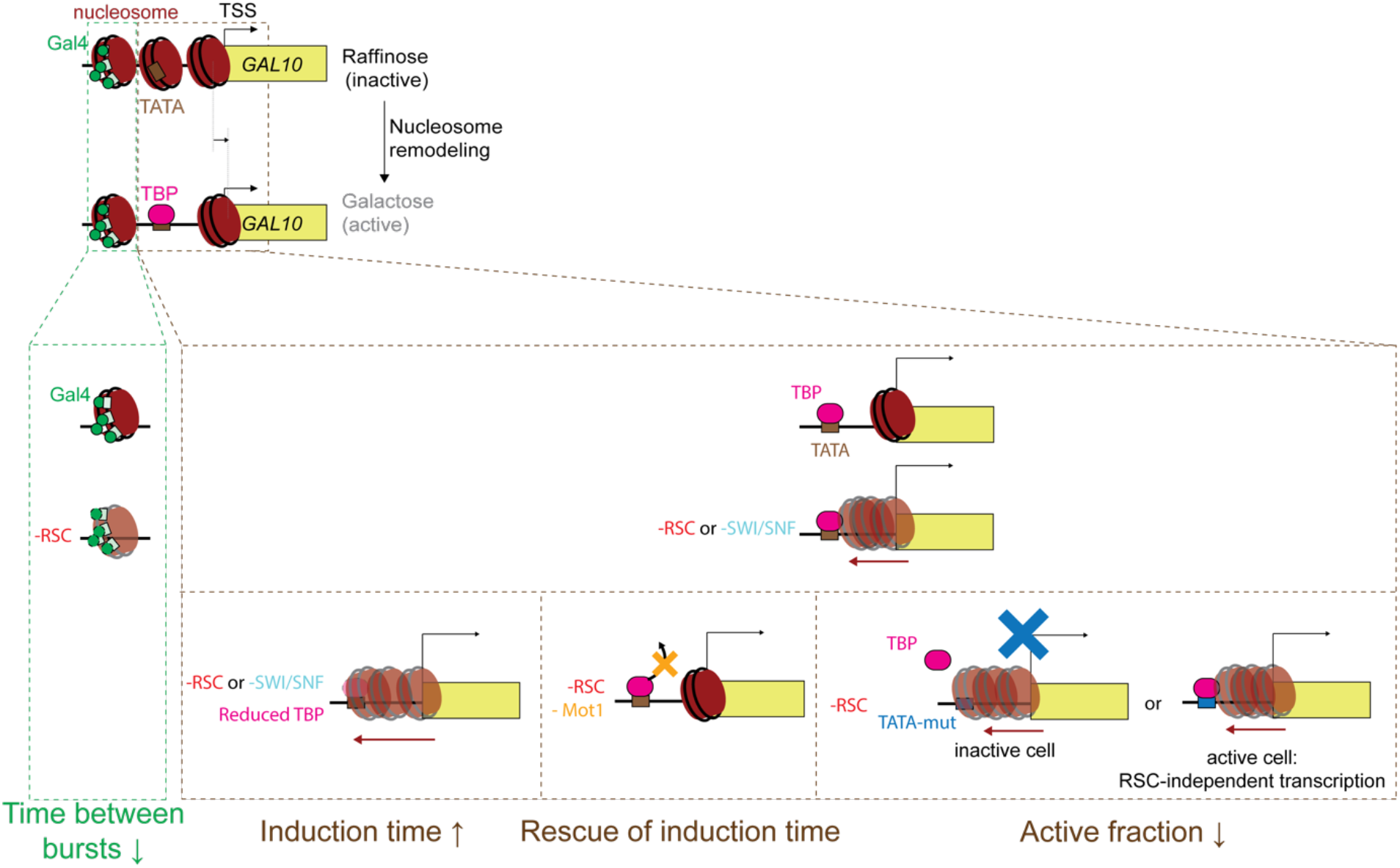
Model showing how remodeling of different nucleosomes regulates transcriptional bursting at the *GAL10* gene. Upon induction of the *GAL10* gene, the three promoter nucleosomes are remodeled to allow for transcription activation. Remodeling of the fragile nucleosome at the Gal4 UASs by RSC and Gal4 binding synergistically regulates time between bursts. The nucleosomes in the region spanning the TATA element and the TSS are redundantly displaced by RSC, SWI/SNF and TBP to regulate the first burst of transcription. Nucleosomes that shift into the promoter upon loss of RSC and/or SWI/SNF are competed away by TBP. In the absence of remodeling by RSC or SWI/SNF, partial depletion of TBP reduces the ability of TBP to compete with nucleosomes, such that induction is synergistically slowed down with RSC depletion. Stabilization of TBP by Mot1 depletion results in increased nucleosome competition and rescue of induction time. Mutation of the canonical TATA box abolished the ability of TBP to compete with nucleosomes, resulting in two populations, where either the nucleosome covered the TATA element resulting in no activation, or the nucleosome was not covering the TATA element resulting in RSC-independent transcription activation.

### Remodeling of the fragile nucleosome at the Gal4 UASs

In line with the role of nucleosomes in forming a barrier for binding of regulatory factors to DNA (Zhu et al., 2018), we find that remodeling of the fragile nucleosome at the Gal4 UAS by RSC and Gal4 binding is required to allow each burst of transcription to start. Similarly, at the CUP1 array, continuous nucleosome remodeling is required to keep the UAS accessible for Ace1 binding at each burst (Mehta et al., 2018). The synergistic effect of reduced Gal4 concentration and RSC depletion indicates that Gal4 binding itself can partially substitute for RSC in configuring the nucleosome to a state which allows transcription activation. A similar destabilization of fragile nucleosome has been observed for general regulatory pioneer factors (GRFs), such as Reb1 or Abf1, although at most genes these GRFs act independently of RSC rather than redundantly like we observe for Gal4 (Kubik et al., 2015, 2018). Mechanistically, Gal4 may trap the nucleosome in a partially unwrapped state, as was shown *in vitro* (Luo et al., 2014). Binding of a gene-specific TF and RSC remodeling thus cooperate to enable efficient TF binding at each burst of transcription.

### Nucleosome remodeling and regulation of TF residence times

We found that RSC depletion and simultaneous depletions of RSC and SWI/SNF modestly reduces burst duration, although only during the first induction and not during re-induction (Figure S3A-E,H-J,S4). Previous work from our lab indicates that the duration of transcriptional bursts is determined by the residence time of Gal4 (Donovan et al., 2019a). The reduced burst duration of the RSC single and RSC&SWI/SNF double depletions found here suggests that these remodelers increase Gal4 residence time. *In vitro*, Gal4 dwell time is considerably reduced by nucleosomes (Luo et al., 2014), and our results suggest that remodelers may counteract this nucleosome-mediated destabilization mechanism *in vivo*. Previous studies have reported effects of RSC on the residence time of different TFs, although in opposing directions. Upon addition of RSC, Rap1 residence time increases *in vitro* as nucleosomes get displaced (Mivelaz et al., 2020), whereas the residence time of Ace1 to the CUP1 array in live yeast decreases in the presence of RSC (Mehta et al., 2018). These contradicting results suggest that the mechanism of TF residence time regulation by RSC remodeling may differ per TF, possibly depending on the ability of the TF to bind to nucleosomal DNA or to displace nucleosomes. The reduced burst duration upon RSC depletion points to a role for RSC in stabilizing Gal4 binding at the fragile nucleosome, but future work is required to directly measure Gal4 residence time *in vivo*, as burst duration may also be affected by the residence time of other transcriptional regulators.

### Nucleosomes at the TATA box

Our study uncovers a novel role for TBP in competing with nucleosomes covering the TATA element. Although TBP depletion in the presence of RSC has a minor effect on nucleosome positioning only at highly expressed genes (Kubik et al., 2018; Tramantano et al., 2016), we show that the effect of TBP on nucleosome displacement becomes more prominent in conditions where nucleosomes cover the TATA box, such as after depletion of RSC or SWI/SNF. Specifically, simultaneous depletion of TBP and RSC results in a larger increase in the nucleosome density around the *GAL10* TATA than single RSC depletion (Figure 4F). This role for TBP in nucleosome positioning appears more important for genes with long TBP residence times and a canonical TATA box (Figure 4,S6). The ability of TBP to compete with nucleosomes allows for remodeler-independent *GAL10* promoter activation, explaining why a significant level *GAL10* of transcription is still observed in the absence of RSC and SWI/SNF.

It was recently shown that TBP is able to bind stably to nucleosomal DNA (Wang et al., 2021). However, in this nucleosomal configuration, TBP cannot efficiently recruit the PIC and activate transcription (Ryan et al., 2000; Wang et al., 2021). Our results reveal that the ability of TBP to compete with nucleosomes depends on the residence time of TBP at the TATA box. We propose a passive competition mechanism where binding of TBP perturbs the stability of the nucleosome and, if the residence time of TBP is long enough, this eventually results in nucleosome eviction or movement of the nucleosome upstream, resulting in a nucleosome-free TATA box needed for recruitment of downstream PIC components. Mutation of the TATA element to reduce TBP residence time results in loss of the ability of TBP to compete with nucleosomes at the TATA element (Figure 5B,C,D,E). Since TBP binding to the TATA box is one of the most stable interactions within the transcription assembly, with a residence time of several minutes (Heiss et al., 2019; Nguyen et al., 2021), we envision that, once bound, TBP can maintain a nucleosome-free TATA element, facilitating initiation of multiple consecutive bursts. TBP binding is thus only rate-limiting for the first burst of transcription, in line with our observed synergy between TBP and RSC in controlling induction time but not time between consecutive bursts.

### Stabilizing TBP binding

To increase the residence time of TBP binding at the *GAL10* promoter and increase the capability of TBP to compete with nucleosomes at the TATA element, we depleted Mot1, the enzyme that clears TBP from TATA and TATA-like sites (Heiss et al., 2019) and found that Mot1 depletion rescues the effect of RSC depletion on *GAL10* induction time. Mot1 is responsible for distribution of TBP over both TATA and TATA-less promoters by removing incorrectly oriented TBP and transcriptionally inactive TBP as well as preventing antisense transcription (Heiss et al., 2019; Koster and Timmers, 2015; Zentner and Henikoff, 2013). In principle, Mot1 depletion could lead to increased antisense transcription and may therefore affect transcription dynamics through a mechanism independent of TBP residence time. However, since we previously showed that the *GAL10* antisense transcript does not affect bursting at *GAL10* (Lenstra et al., 2015), it is unlikely that the rescue of RSC depletion phenotype by Mot1 is indirectly caused by antisense transcription. We instead propose that upon Mot1 depletion an increase residence time of TBP binding at TATA promoters (Spedale et al., 2012; Xue et al., 2017; Zentner and Henikoff, 2013) is sufficient to achieve an active nucleosome configuration around the TATA element, such that RSC remodeling is no longer needed.

### Taf1 binding around the +1 nucleosome

The observed nucleosome competition is specific for TBP, rather than being a common mode of action of all PIC components, since Taf1 depletion does not show the same synergy with RSC depletion as TBP depletion. Even though Taf1 interacts stably with the +1 nucleosome in yeast extracts (Joo et al., 2017), Taf1, as well as other PIC components, interacts with chromatin only for a few seconds *in vivo* (Nguyen et al., 2021), contradicting the stable chromatin engagement that may be needed to passively compete away nucleosomes such as we propose for TBP (Mivelaz et al., 2020). In addition, upon Taf1 depletion, we observe no effect on burst duration, suggesting that the TF-independent reinitiation mechanism observed *in vitro* (Joo et al., 2017) may be less important for the TF-driven activation at *GAL10*. Moreover, we observe a lower induction time than expected for simultaneous depletion of Taf1 and RSC, indicating that Taf1 and RSC have opposing functions. In support, recent single-molecule tracking measurements of PIC components revealed that Taf1 binding to chromatin becomes more stable upon RSC depletion, suggesting that RSC promotes TFIID turnover (Nguyen et al., 2021). Our data thus indicates that RSC, SWI/SNF and TBP redundantly are able to displace nucleosomes around the TATA box and that RSC inhibits stable Taf1 binding around the TSS and +1 nucleosome.

Overall, dynamic epistasis analysis provides detailed mechanistic insight into how nucleosome remodeling acts in combination with transcription factors and assembly of the PIC to control the kinetics of transcriptional bursting. In particular, at the yeast *GAL10* gene and other TATA-containing genes, a role for TBP in competing with nucleosomes *in vivo* is uncovered, which together with chromatin remodelers enables efficient PIC assembly and transcription initiation.

## Acknowledgements

We thank the Holstege laboratory for strains. We thank Aleksandra Balwierz, Linda Joosen, Judith Haarhuis, Hans Teunissen, Laureen Willems, Robin van der Weide for assistance with MNase-seq experiments. We thank the Research High Performance Computing Facility and the Genomics Core Facility of the NKI for assistance. We thank members of the T.L.L. and F.v.L. laboratories for helpful discussions and Elzo de Wit and members of the T.L.L. laboratory for critical reading of the manuscript. This work was supported by the Netherlands Organization for Scientific Research (NWO, 016.Veni.192.071 (I.B.), ZonMW-TOP 91218022 (F.v.L.) and gravitation program CancerGenomiCs.nl (T.L.L.)), Oncode Institute (T.L.L.), which is partly financed by the Dutch Cancer Society, and the European Research Council (ERC Starting Grant 755695 BURSTREG (T.L.L.)).

## Author contributions (CRediT taxonomy)

Conceptualization: I.B., T.L.L.; Methodology: I.B.; Software: I.B., T.L.L.; Formal analysis: I.B.; Investigation: I.B., E.K.; Writing—Original draft: I.B., T.L.L.; Writing—Review and Editing: F.v.L; Supervision; F.v.L, T.L.L.; Funding Acquisition: I.B., F.v.L, T.L.L.

## Declaration of interests

The authors declare no competing interests

## STAR methods

### Yeast strains and plasmids

All strains were derived from BY4741 and BY4742 anchor-away background strains (Jonge et al., 2017). The relevant FRB-tags for anchor-away were introduced either by transformation with a PCR product containing the FRB-yEGFP1-hphMX4 cassette (pTL100) or the FRB-mScarlet-hphMX4 cassette (pTL329). Alternatively, a CRISPR-Cas9-based approach was used (Laughery et al., 2015), and strains were transformed using a plasmid expressing Cas9, a guide RNA and double-stranded PCR repair template from the same plasmids, followed by removal of the Cas9 plasmid by 5-FOA selection. PP7 loops were introduced by transformation with a PCR product containing the PP7 loop cassette and loxP-kanMX-loxP (pTL031). The kanMX marker was removed by expressing CRE recombinase (pTL014 or pTL191). PP7-coat protein was integrated at the URA3 locus by transformation of PacI-digested single-integration plasmid (pTL174) (Wosika et al., 2016). TATA-mut-int2 (Blake et al., 2006) was introduced in TATA element of *GAL10* by the CRISPR-Cas9 approach described above using a single-stranded oligo and a plasmid expressing Cas9 and a guide RNA. The *GAL4/gal4Δ* strain was created by mating a BY4741 anchor-away strain with wildtype *GAL4* with a BY4742 anchor-away *gal4Δ* haploid strain that was constructed by the CRISPR-Cas9 approach described above using a single-stranded oligo and a plasmid expressing Cas9 and a guide RNA. For all strains, at least 2 replicates were constructed independently, which were verified by PCR, growth plates (Figure S2A), microscopy (Figure S2B) and if applicable, sequencing and smFISH (Figure S2C). All strains, plasmids and oligos used in this study are listed in Table S1, S2 and S3 respectively.

### Live-cell imaging of transcription dynamics

Live-cell imaging of transcription dynamics was performed as previously described in detail in (Brouwer et al., 2020; Donovan et al., 2019a) with minor modifications. In brief, cells were treated with 7.5 µM rapamycin or DMSO for 60 min and subsequently imaged at mid-log (OD600_nm_ 0.2-0.4) on a coverslip with an agarose pad consisting of 2% agarose and synthetic complete medium containing 2% galactose and 7.5 µM DMSO or rapamycin. Imaging was performed on a setup consisting of an AxioObserver inverted microscope (Zeiss), an alpha Plan-Apochromat 100x NA 1.46 oil objective, an sCMOS ORCA Flash 4v3 (Hamamatsu) with a 475-570 nm dichroic (Chroma), 570 nm longpass beamsplitter (Chroma) and 515/30 nm emission filter (Semrock), a UNO Top stage incubator (OKOlab) at 30 C, and LED excitation at 470/24 nm (SpectraX, Lumencor) at 20% power and an ND 2.0 filter, resulting in a 62 mW/cm^2^ excitation intensity. Widefield images were recorded 1 hour at 15 s interval, with *z*-stacks (9 slices, *Δ*z 0.5 µm) and 150 ms exposure using Micro-Manager software (Edelstein et al., 2014). For each condition, at least 3 replicate datasets were acquired with in total at least 100 cells.

### Microscopy of anchor away nuclear depletion

Before acquiring transcription dynamics images, proper nuclear depletion by anchor away (Haruki et al., 2008) was ensured in each sample by imaging of FRB-mScarlet tag. Imaging was performed on the setup described above but with 475-570 nm dichroic (Chroma), 570 nm longpass beamsplitter (Chroma) and 600/52 nm emission filter (Semrock), and LED excitation at 550/15 nm (SpectraX, Lumencor) at 100% power and ND2.0 filter, resulting in 413.0 mW/cm^2^ excitation intensity. A single widefield image was recorded as a *z*-stack (9 slices, *Δz* 0.5 µm) using either 150 ms exposure (for YTL1178, YTL1179, YTL1281, YTL1309) or 500 ms exposure (for YTL1505, YTL1506, YTL1508, YTL1510, YTL1448, YTL1450, YTL1470, YTL1591, YYL1626).

### Analysis of transcription dynamics

For analysis of the transcription dynamics imaging data, a similar approach was used as described in (Donovan et al., 2019a). All analysis was implemented as custom-written Python software (https://github.com/Lenstralab/livecell_analysis). First, images were corrected for *xy*-drift in the stage using an affine transformation on the maximum intensity projection. Next, cells were segmented using Otsu thresholding and watershedding. The intensity of the TS was calculated by fitting a 2D Gaussian mask after local background subtractions as described previously (Coulon et al., 2014). Initially, a threshold of 6 times the standard deviation of the background was used. For frames where no TS was detected, a second fit was made in the vicinity of the high intensity spots detected in that cell, using a threshold of 4 times the standard deviation of the background. For frames where no TS was detected after this second fit, the intensity was measured at the location of the previous frame where a TS was successfully found. The tracking within each cell was inspected visually, and the endpoint of each trace was manually set at the last frame where a TS is visible. Cells in which a TS was not reliably detected were excluded from the analysis.

To determine the on and off periods, binarization was performed using a threshold set at 5 times the standard deviation of the background. The standard deviation of the background was determined for each cell fitting by a Lorentzian distribution to intensities measured at four points at a fixed distance from the TS in each frame in the same cell. This threshold was chosen to reliably distinguish on and off periods from background levels at the single-transcript level. Subsequently, the binarization was improved by removing bursts that last a single frame and merging bursts that are separated by a single frame. From these binarized traces, the burst durations, time between bursts, induction time are directly calculated. The burst intensity is measured as the average intensity of all frames in which the cell was on. The fraction of active cells was determined by manual scoring of the cell that do and the cells that do not show a TS during the 1-hour acquisition.

In total at least 100 cells were included for each condition, and values for burst duration, time between bursts, induction time and burst intensity are determined by bootstrapping with 1000 repetitions. Reported error bars are standard deviations from the same bootstrap. Error bars in the number of active and inactive cells are given by the square root of the number of cells, as cells are expected to be independent of each other and thus follow Poisson statistics. To determine whether the obtained bursting parameters are significantly different between conditions, we have used bootstrap hypothesis testing using equation (4) from (MacKinnon, 2007) to determine the achieved significance level.

### Dynamic epistasis analysis

For the dynamic epistasis analysis we devised in this study, the fractional change in each parameter of transcriptional bursting was determined as the ratio between the bootstrap mean of this parameter in the perturbed population and the unperturbed population. Error bars were calculated from the same bootstraps and propagated under the assumption that the measurements are independent between conditions. To calculate the expected effect of a double perturbation on each parameter, fractional changes of the individual perturbations are multiplied, analogous to to the way phenotypic growth effects caused by pairwise genetic interactions are assessed (van Leeuwen et al., 2017). Error bars are calculated by error propagation of the errors of individual perturbations.

### Fitting of induction time distributions

To determine whether gene induction depends on a single or multiple rate-limiting steps, a least-squares fit was performed on the histogram of the distribution of induction times with a binsize of 1 minute. The following parameterization of the probability density function of the Gamma distribution was used:

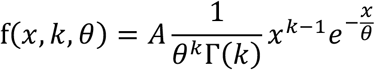

Here, Γ(*k*) is the Gamma function, defined as: 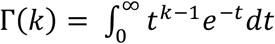

The amplitude parameter *A* was added because there is a dead-time between addition of galactose and actual start of the image acquisition. Therefore, a varying fraction of the full distribution is observed and normalization to 1 would be inappropriate. Free parameters in the fit are *A, k* and *θ*. For the parameter *A*, the lower bound is set at 1 and the initial guess is 10. For *k*, the lower bound is set at 1.0001 and the initial guess is set at 10 (for non-memory induction) or at 1.0001 (for all reinduction conditions). For *θ*, the lower bound is set at 0.0001 and the initial guess at 1. The scale parameter *k* is a measure for the number of rate-limiting steps. Therefore, we interpret distributions with a value of *k* that is not different from 1 as governed by a single rate-limiting step and distributions with a value of *k* that is larger than one as being governed by multiple rate-limiting steps.

### Single-molecule RNA FISH

Single-molecule RNA fluorescence in situ hybridization (smFISH) was performed as previously described (Donovan et al., 2019a; Patel et al., 2021) with minor modifications. In brief, yeast cultures were grown to early mid-log (OD_600nm_ 0.5), treated with either 7.5 µM rapamycin or DMSO for 60 min for anchor-away prior to fixation with 5% PFA (Electron Microscopy Sciences, 15714-S) for 20 min. Then cells are washed three times with buffer B (1.2 M sorbitol and 100 mM potassium phosphate buffer pH 7.5), permeabilized with 300 U of lyticase (Sigma-Aldrich, L2524-25KU), and washed with buffer B. Cells were immobilized on poly-L-lysine-coated coverslips (Neuvitro) and permeabilized with 70% ethanol overnight or up to 3 days. Coverslips were hybridized for 4h at 37°C with hybridization buffer containing 10% dextran sulfate, 10% formamide, 2xSSC, and 5 pmole probe. Four PP7 probes labeled with Cy3 (for YTL1178, YTL1179, YTL1281 and YTL1309) or Cy5 (for YTL1448, YTL1450, YTL1505, YTL1506, YTL1508, YTL1510, YTL1470, YTL1591, YTL1626) were targeted to the loops, or 48 probes labeled with Quasar670 (for YTL524, YTL525, YTL526, YTL527, YTL528, YTL529) were targeted to coding region of *GAL10* (Table S4). Coverslips were washed 2× for 30 min with 10% formamide, 2xSSC at 37°C, 1× with 2xSSC, and 1× for 5 min with 1× PBS at room temperature. Coverslips were mounted on microscope slides using ProLong Gold mounting media with DAPI (Thermo Fisher, P36934).

Imaging was performed on two similar microscope setups consisting of an AxioObserver inverted microscope (Zeiss), a Plan-Apochromat 40x NA 1.4 oil DIC UV objective, a 1.25x optovar, an sCMOS ORCA Flash 4v3 (Hamamatsu). For Cy3, we used a 562 nm longpass dichroic (Chroma), 595/50 nm emission filter (Chroma) and 550/15 nm LED excitation at full power (Spectra X, Lumencor), with an excitation intensity at the two microscopes of 6.8 W/cm^2^ or 8.8 W/cm^2^. For Cy5, we used a 660 nm longpass dichroic (Semrock or Chroma), 697/60 nm emission filter (Chroma) and 640/30 nm LED excitation at full power (Spectra X, Lumencor), with an excitation intensity at the two microscopes of 4.9 W/cm^2^ or 6.7 W/cm^2^. For DAPI, we used either a 410nm/490nm/570nm/660nm dichroic (Chroma), a 430/35 nm, 512/45 nm, 593/40 nm, 665 nm longpass emission filter (Chroma) or a 425 nm longpass dichroic (Chroma) and a 460/50 nm emission filter (Chroma) and LED excitation at 395/25 nm at 25% power (Spectra X, Lumencor), with an excitation intensity at the two microscopes of either 1.2 W/cm^2^ or 1.9 W/cm^2^. For each sample, for at least 50 fields-of-view, a *z*-stack (21 slices, *Δz* 0.3 µm) was recorded for DAPI and Cy3 or Cy5, using 25 ms exposure for DAPI and 250 ms exposure for Cy3 and Cy5 using Micro-Manager software.

### Analysis of smFISH

Images were analyzed using custom-written Python software (https://github.com/Lenstralab/smFISH_analysis). Here, cells and nuclei were segmented using Otsu thresholding and watershedding. Spots were localized by fitting a 3D Gaussian mask after local background subtraction (Coulon et al., 2014). Cells in which no spots were detected were excluded from further analysis, because visual inspection indicated these cells were not properly segmented or not properly permeabilized such that smFISH probes did not enter the cells. For each cell, the TS was defined as the brightest nuclear spot and the number of RNAs at each TS was determined by normalizing the intensity of each TS with the median fluorescent intensity of the cytoplasmic RNAs detected in all cells. Cells with fewer than 5 RNAs at the TS were classified as inactive, and cells with 5 or more RNAs at the TS were classified as active cells. Subsequently, the fraction of active cells and the mean number of RNAs at the TSs of active cells were determined. For each condition, at least 3 replicate experiments were performed with in total at least 5000 cells, and the average value and standard error of the mean were determined for both the active fraction and the number of RNAs at the TSs of active cells. The fractional changes of these parameters upon nuclear depletion of indicated factors were determined from these mean values.

For classification of cells into G1, S and G2 cell-cycle stage, the sum of the nuclear DAPI intensity in each cell is calculated from a maximum intensity projection. Subsequently, a histogram of all nuclear DAPI intensities (with 50 equally spaced bins) is fit with a Gaussian mixture model consisting of 2 peaks. Cells are classified as G1 stage if they are in a window of (SD_1_, 0.75xSD_1_) around the center of the first peak, as G2 stage if they are in a window of (0.5xSD_2_, 1.5xSD_2_) around the center of the second peak and as S stage if they are in between the 2 peaks, where SD_1_ and SD_2_ are the standard deviations of the first and second peak, respectively. Fractional changes in active fraction and number of RNAs at the TSs of active cells are determined as described above for each cell-cycle stage separately.

### MNase-seq

Preparation and analysis of mono-nucleosomal DNA was performed as described previously (Donovan et al., 2019a; Jonge et al., 2017) with minor modifications. Briefly, cells were grown in SC + 2% raffinose or SC + 2% galactose from OD 0.3 to OD 0.75 and then treated with 7.5 µM rapamycin or DMSO for 60 min. Then, cells were fixed in 1% PFA, washed with 1 M sorbitol, treated with spheroplasting buffer (1M sorbitol, 1 mM β-mercaptoethanol, 10 mg/ml zymolyase 100T (US biological, Z1004.250)) and washed twice with 1 M sorbitol. Spheroplasted cells were treated with 0.01171875 U (low MNase) or 0.1875 U (high MNase) micrococcal nuclease (Sigma-Aldrich, N5386-200UN) in digestion buffer (1 M sorbitol, 50 mM NaCl, 10 mM Tris pH7.4, 5 mM MgCl2, 0.075% NP-40, 1 mM β-mercaptoethanol, 0.5 mM spermidine) at 37°C. After 45 min, reactions were terminated on ice with 25 mM EDTA and 0.5% SDS. Samples were treated with proteinase K for 1 h at 37°C and decrosslinked overnight at 65°C. Digested DNA was extracted with phenol/chloroform (PCI 15:14:1), precipitated with NH_4_-Ac, and treated with 0.1 mg/ml RNaseA/T1. The extent of digestion was checked on a 3% agarose gel. For all conditions, two independent experiments were performed, with similar outcomes, except for the SWI/SNF-depletion strain treated with high MNase concentration, where one replicate of the DMSO condition was underdigested. For this condition, only one replicate was used for analysis.

Sequencing libraries were prepared using the KAPA HTP Library Preparation Kit (07961901001, KAPA Biosystems) using 1 µg of input DNA, 5 µL of 10 µM adapter, double-sided size selection before and after amplification using 10 cycles. Adapters were created by ligation of Universal adapter to individual sequencing adapters (Table S5). Libraries were checked on Bioanalyzer High Sensitivity DNA kit (agilent). Sequencing was performed on a NextSeq550. Paired-end 2×75 bp reads were aligned to the reference genome SacCer3 (January 2015) using bowtie2 (Langmead and Salzberg, 2012) with the settings *--sensitive -- end-to-end -3 15 -5 5 -X 1980 --no-contain --no-discordant -p 40 -x*. The data have been deposited in NCBI’s Gene Expression Omnibus (Edgar et al., 2002) and are accessible through GEO Series accession number GSE190737.

### Analysis of MNase-seq

Analysis of MNase-seq data was done using custom-written Python software (https://github.com/Lenstralab/MNase_analysis). First, the aligned reads were filtered for length and only reads between 95 and 225 bp were retained for analysis. Subsequently, the read coverage was determined on a chromosome-by-chromosome basis by counting the number of reads covering each base, and the normalized coverage was determined by normalizing the read coverage to the total coverage on the chromosome. Next, the normalized coverage along each gene was determined using all open reading frames that are classified as ‘verified’ in the Saccharomyces Genome Database (SGD) (Cherry et al., 2012). TATA and TATA-mismatch genes were identified as ‘TATA-containing’ and ‘TATA-less’ from (Rhee and Pugh, 2012). The coverage in TATA or TATA-mismatch regions was determined as the sum of the normalized coverage in the 8-bp region spanning the TATA or TATA-mismatch sequence as determined by (Rhee and Pugh, 2012).

For metagene plots, genes were aligned at the +1 nucleosome in unperturbed conditions. To determine the position of the +1 nucleosome for each gene in these unperturbed conditions, the (pre-normalization) coverage in a 4000 bp window around the TSS of each gene was extracted from all experiments performed in DMSO using the high MNase concentration (combining the data for all yeast strains, i.e. YTL524, YTL525, YTL1306 and YTL1584). If the gene was on the Crick-strand, the coverage was flipped to facilitate alignment of all genes. For each gene, these coverages were subsequently summed and smoothed using a Gaussian filter with a 40 bp window. The minimum of this smoothed coverage was determined, and a peak-calling function was used to detect nucleosome peaks. The -1 nucleosome was defined as the first peak before the coverage minimum, and the +1 nucleosome as the first peak after the coverage minimum. Genes for which fewer than 2 peaks were detected or for which the -1 or +1 nucleosome was detected more than 1000 bp away from the TSS were excluded from the analysis. To generate metagene plots, the normalized coverage of all genes in a window of 2000 bp centered at the location of the +1 nucleosome in unperturbed conditions of that gene was averaged. To generate heatmaps of the log2-fold-change of the coverage upon depletions, genes were sorted by the NDR width as determined by the distance between the -1 and +1 nucleosomes in unperturbed conditions. Subsequently, for each gene, the log2-fold-change between the coverage in each depletion (rapamycin) condition and the average coverage between two replicate experiments in non-depleted (DMSO) condition was calculated. This data was represented as a heatmap. To determine the shift in +1 position, the location of the +1 nucleosome was determined in each depletion dataset independently using the same steps as performed on the summed coverage to detect the position of the +1 nucleosome in unperturbed conditions. The shift in +1 nucleosome was then defined as the difference between the +1 nucleosome in depleted conditions and the +1 nucleosome as determined from all unperturbed high MNase datasets.

### Growth assay

The growth assay used to assess growth rapamycin-or DMSO-containing plates was performed as described previously (Donovan et al., 2019a) with minor modifications. Serial five-fold dilutions of (YTL559, YTL658, YTL047, YTL525, YTL524, YTL1306, YTL1281, YTL1391, YTL1413, YTL014, YTL1397, YTL1506, YTL1584, YTL1510, YTL1613, YTL1615, YTL1394, YTL1588) strains were spotted on YEP + 2% glucose + 7.5 µM rapamycin, YEP + 2% glucose + DMSO, YEP + 2% galactose + 20 μg/μl ethidium bromide + 7.5 µM rapamycin, YEP + 2% galactose + 20 μg/μl ethidium bromide + DMSO, and YEP + 2% raffinose + 2% galactose + 40 mM lithium chloride + 0.003% methionine + 7.5 µM rapamycin amd YEP + 2% raffinose + 2% galactose + 40 mM lithium chloride + 0.003% methionine + DMSO. Growth was assessed after 3 days at 30°C.

## Notes

https://www.ncbi.nlm.nih.gov/geo/query/acc.cgi?acc=GSE190737

### Competing Interest Statement

The authors have declared no competing interest.

