## Supplemental Information for "Dynamic epistasis analysis reveals how chromatin remodeling regulates transcriptional bursting"

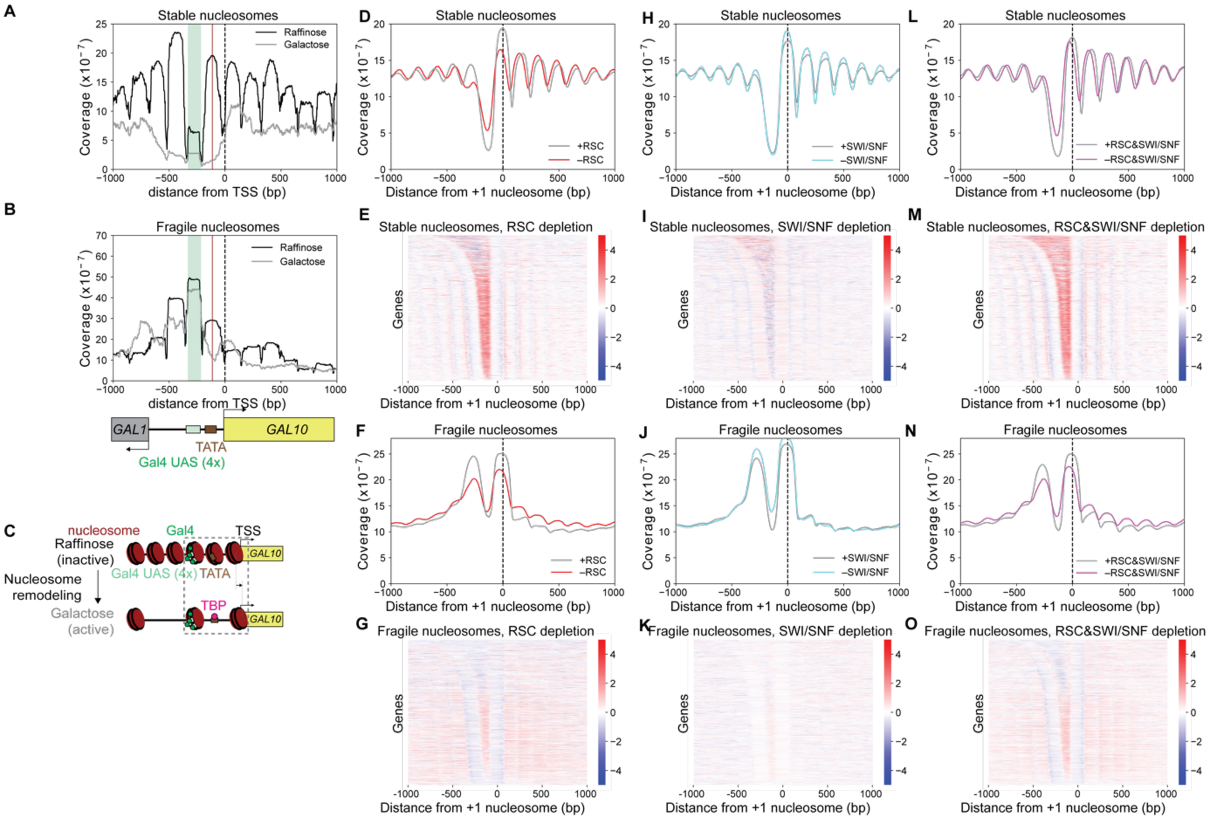


**Figure S1, related to Figure 1 and Figure 2**

**Genome-wide changes in nucleosome coverage upon depletion of RSC, SWI/SNF and RSC&SWI/SNF.**

(A),(B) MNase-seq analysis of (A) stable and (B) fragile nucleosomes in the *GAL10* promoter region in active (galactose) and inactive (raffinose) conditions.

(C) Schematic representation of nucleosome remodeling during activation of *GAL10.* Grey box: region with nucleosomes important for regulation of *GAL10* that are discussed in this study.

(D),(F),(H),(J),(L),(N) Metagene MNase-seq analysis of stable or fragile nucleosomes (as indicated in the figure) upon RSC depletion by anchor-away of Sth1, SWI/SNF depletion by anchor-away of Swi2, or simultaneous depletion of RSC and SWI/SNF, averaged over all genes as annotated by (Cherry et al., 2012), aligned at the location of the +1 nucleosome (black dashed line).

(E),(G),(I),(K),(M),(O) Heatmap of the log2-fold-change in nucleosome coverage of stable or fragile nucleosomes (as indicated in the figure) at all genes as annotated by (Cherry et al., 2012), sorted by NDR width and aligned at the location of the +1 nucleosome, upon depletion of RSC, SWI/SNF or RSC and SWI/SNF depletion.

Shown is one representative replicate out of two experiments for all except (I) and (J), which is a single replicate experiment.

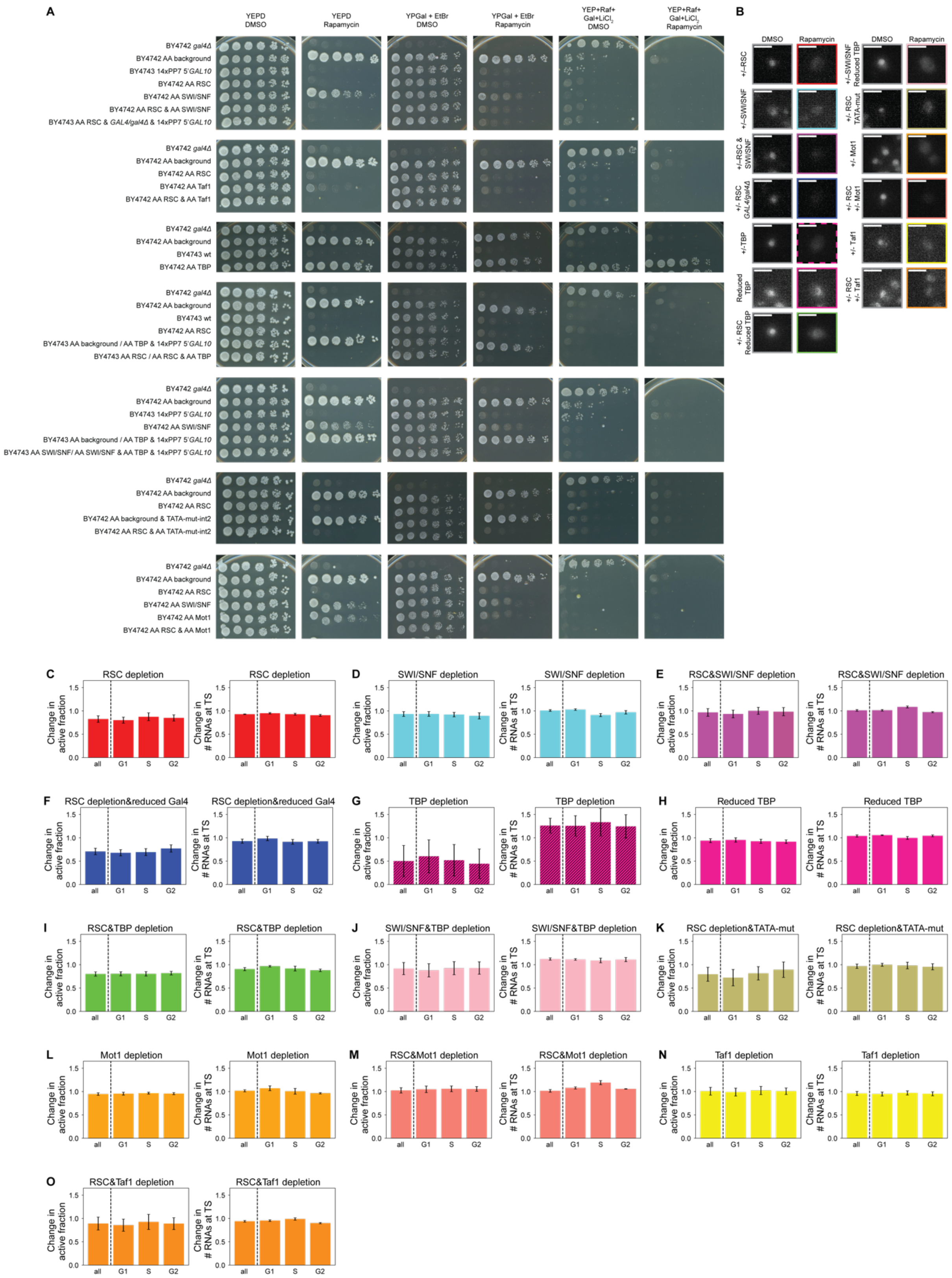


**Figure S2, related to Figure 1, Figure 2, Figure 3, Figure 4, Figure 5 and Figure 6**

**Verification of strains used for nuclear protein depletions using anchor away.**

(A) Growth of anchor-away yeast strains used in this study assessed on YEPD, YEP + Galactose + Etidium bromide and YEP + Galactose + Raffinose + Lithium Chloride plates with either DMSO or rapamycin. Shown are (1:5) serial dilutions of cultures, starting at $OD_{600nm}$ of 0.3.

(B) Depletion of all indicated factors was verified by imaging cells after 60 min of rapamycin-treatment or control cells treated with DMSO. Shown is 1 typical cell per condition, out of at least 100 cells from three biological replicates.

(C-O) No effect of cell-cycle stage on the fraction of (left) active cells and (right) number of RNAs at the TS active cells upon depletion of indicated factors based on smFISH experiments. Active cells defined as cells with 5 or more RNAs at TS. Error bars are SEOM from 3 independent experiments.

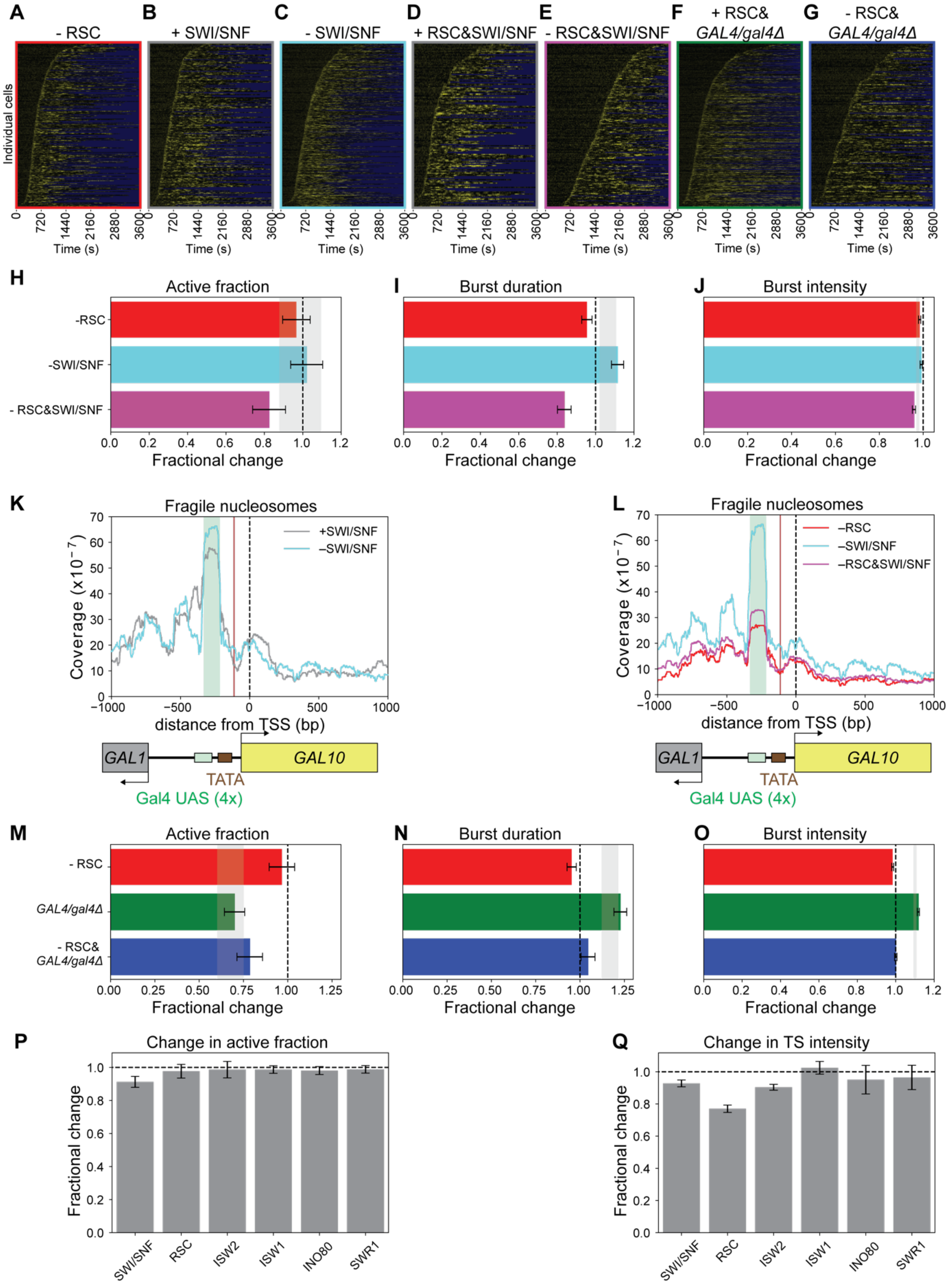


**Figure S3, related to Figure 1, Figure 2 and Figure 3**
**Change in transcription dynamics and nucleosome coverage upon single and double depletion of chromatin remodeling complexes.**

(A-G) Heatmap of the TS intensity in (A) N=292, (B) N=213, (C) N=453, (D) N=151, (E) N=202, (F) N=471, and (G) N=181 cells (rows) in the presence or absence of indicated factors. Yellow: fluorescence intensity of TS; blue: region excluded from analysis.

(H-J) The (H) fraction of cells that activated during 1 hour of imaging, (I) the burst duration and (J) the burst intensity for single and double depletions of RSC and SWI/SNF. Grey bar: expected effect based on dynamic epistasis analysis. Error bars are propagated standard deviations based on 1000 bootstrap repeats.

(K) MNase-seq analysis of fragile nucleosomes in the *GAL10* promoter region showed no change upon depletion of SWI/SNF.

(L) MNase-seq analysis of fragile nucleosomes in the *GAL10* promoter region upon simultaneous depletion of RSC and SWI/SNF compared to depletion of either RSC or SWI/SNF individually.

(M-O) The (M) fraction of cells that activated during 1 hour of imaging, (N) the burst duration and (O) the burst intensity for depletion of RSC, in *GAL4/gal4Δ* cells and the double perturbation . Grey bar: expected effect based on dynamic epistasis analysis. Error bars are propagated standard deviations based on 1000 bootstrap repeats.

(P,Q) Change in fraction of active cells (I) and change in transcription site (TS) intensity of the active cells based on smFISH experiments upon depletion of the catalytic subunits of each of the yeast chromatin remodeling complexes. Active cells defined as cells with fewer than 5 RNAs at the TS. Error bars are SEOM from 3 independent experiments.

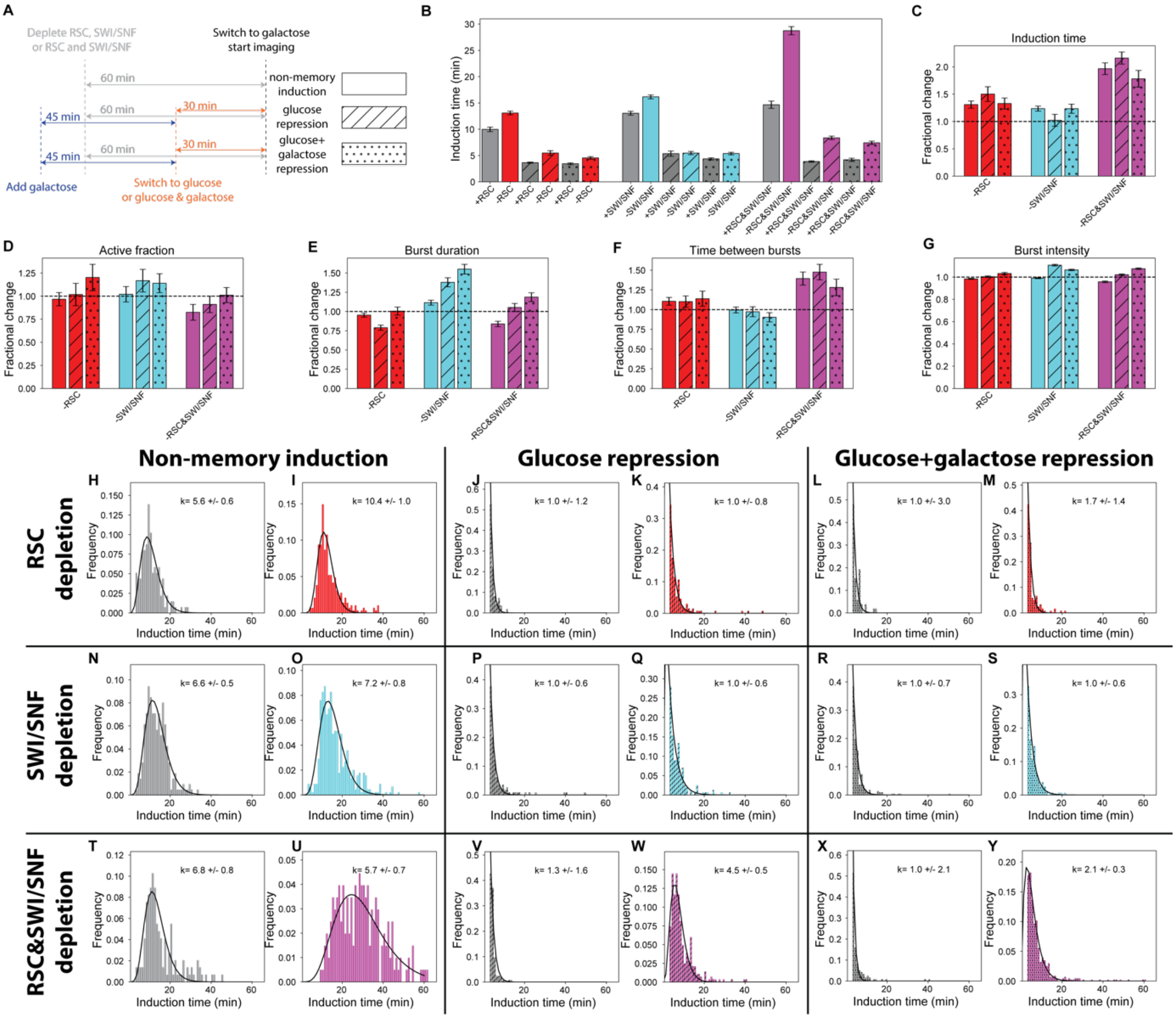


**Figure S4, related to Figure 1 and Figure 2**

**Transcriptional re-induction of *GAL10* shows increase in the number of activation steps upon RSC and SWI/SNF depletion.**

(A) Schematic explaining re-induction experiments. For non-memory induction, cells were grown in raffinose. For memory experiments, cells were primed by adding galactose for 45 min and subsequently washed and resuspended in the appropriate repression sugar 30 min before imaging. As in all experiments, rapamycin or DMSO was added for depletion 60 min prior to galactose addition and imaging.

(B) *GAL10* induction time in memory and non-memory conditions, upon depletion of RSC, SWI/SNF or both RSC and SWI/SNF. Error bars are standard deviations based on 1000 bootstrap repeats.

(C-G) Change in (C) induction time, (D) active fraction, (E) burst duration, (F) time between bursts and (G) burst intensity upon depletion of RSC, SWI/SNF or both RSC and SWI/SNF. Error bars are propagated standard deviations based on 1000 bootstrap repeats.

(H-Y) Distribution of induction time in memory or non-memory conditions as indicated on top in presence (grey) or absence (red, cyan or magenta) of remodeler indicated on the left. Black line: fit with Gamma distribution; inset: shape parameter *k* obtained from fit. A *k*-value significantly different from 1 indicates a single rate-limiting step, while $k > 1$ indicates multiple rate-limiting steps. Without remodeler depletion, re-induction after glucose repression shows a single rate-liming step (V,X), which increases to multiple rate-limiting steps after combined RSC&SWI/SNF depletion (W,Y).

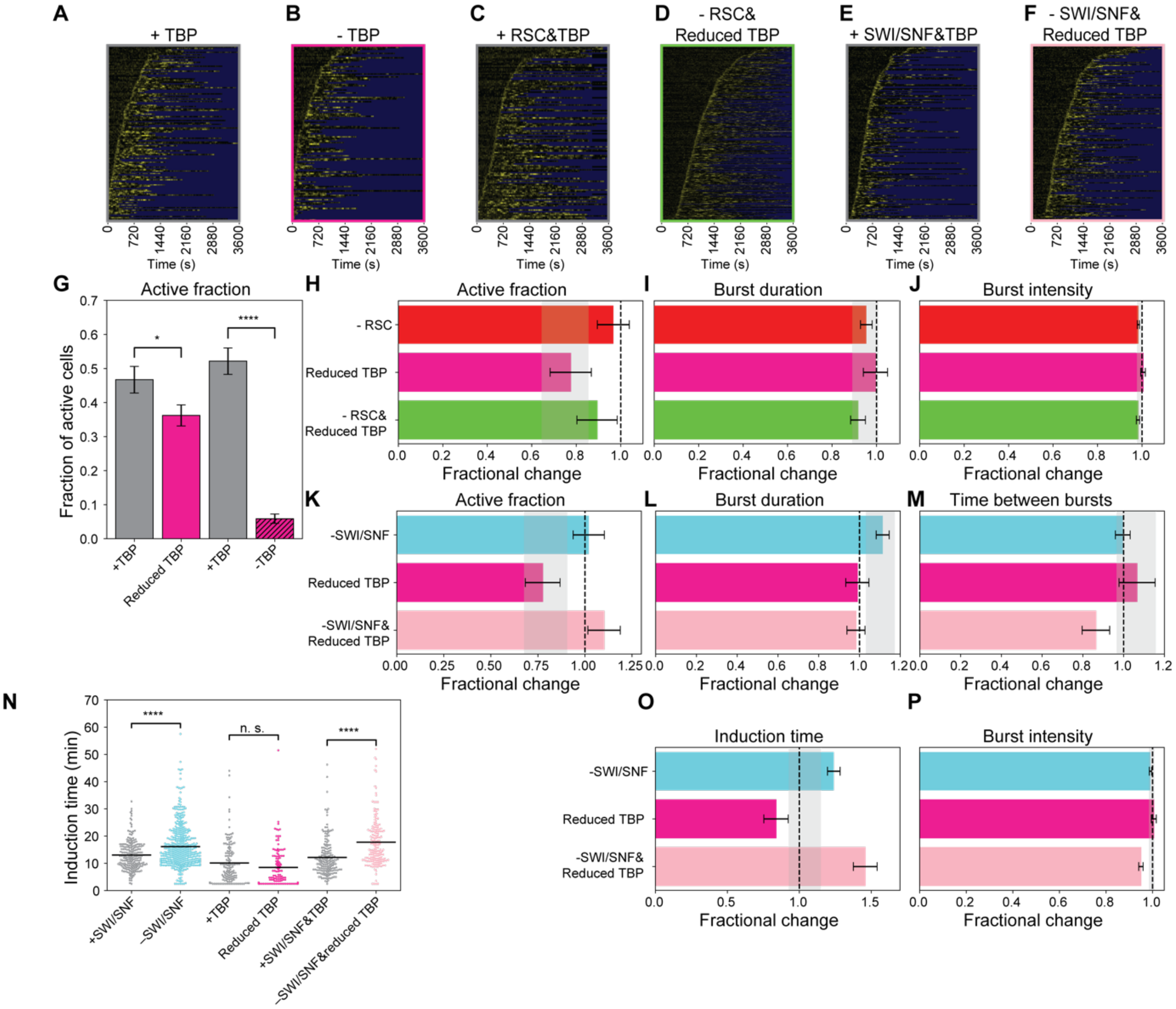


**Figure S5, related to Figure 4**

**Transcription dynamics upon perturbed nucleosomes around the TATA element, by depletion of TBP in combination with RSC or SWI/SNF.**

(A-F) Heatmap of the TS intensity in (A) N=139, (B) N=120, (C) N=122, (D) N=340, (E) N=179 and (F) N=175 cells (rows) in the presence or absence of indicated factors. Yellow: fluorescence intensity of TS; blue: region excluded from analysis.

(G) The fraction of cells that activated during 1 hour of imaging when depleting TBP fully (-TBP, both copies tagged for depletion) or partially (reduced TBP, one of two copies tagged for depletion) using anchor-away.

(H-J) The fraction of (H) cells that activated during 1 hour of imaging, (I) the burst duration and (J) the burst intensity, when simultaneously depleting RSC and reducing TBP levels were as expected based on their individual depletions. Grey bar: expected effect based on dynamic epistasis analysis.

(K-P) SWI/SNF and simultaneous TBP and SWI/SNF depletion showed (K) increased induction time of *GAL10*. (L) The fraction of cells that activated during 1 hour of imaging, (M) the burst duration, (N) the time between bursts, (O) the induction time and (P) the burst intensity for single and double depletion of SWI/SNF and reduced TBP. Grey bar: expected effect based on dynamic epistasis analysis.

Error bars in (A-E) are propagated standard deviations based on 1000 bootstrap repeats. Significance in (A) and (E) determined by bootstrap hypothesis testing (MacKinnon, 2007); n.s.: not significant; ****: $p < 0.00005$.

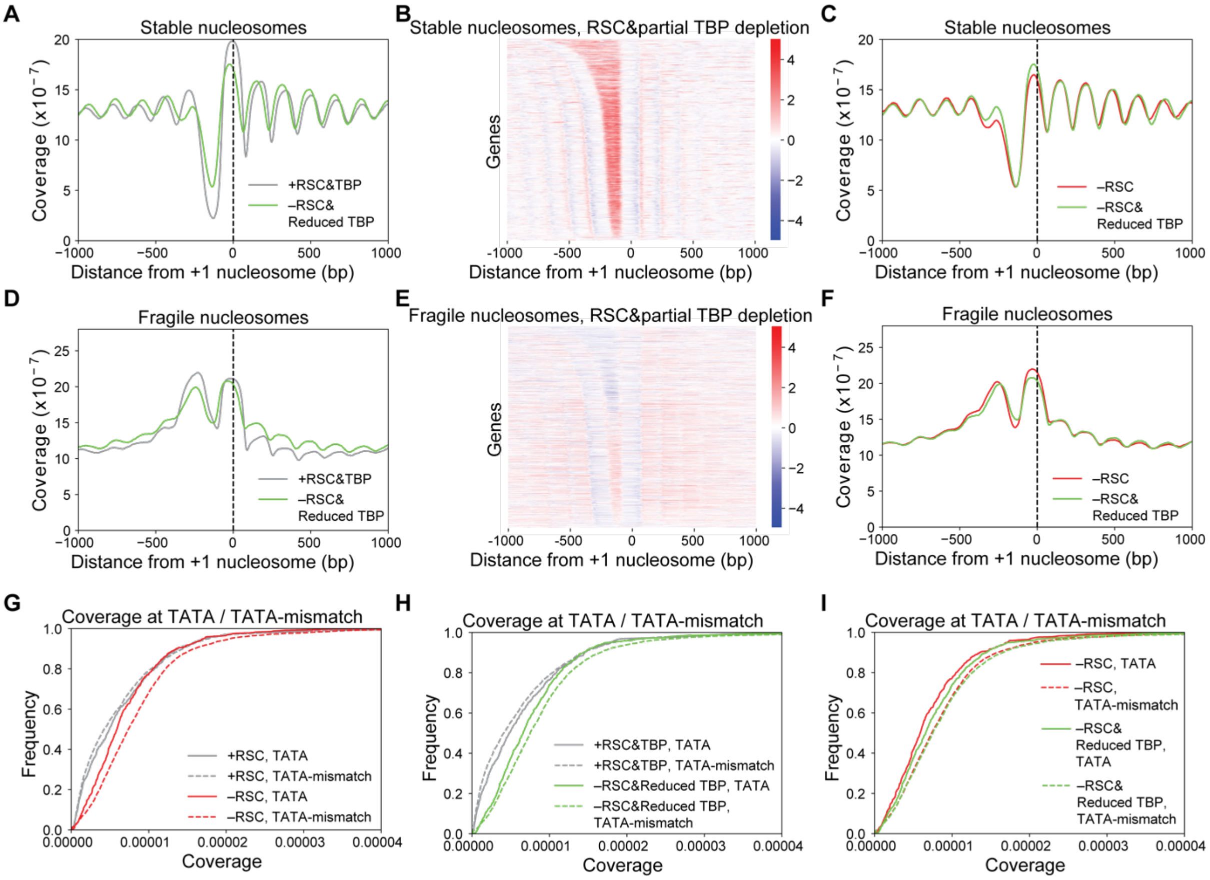


**Figure S6, related to Figure 4**

**Genome-wide changes in nucleosome coverage upon depletion of RSC and TBP.**

(A),(D) Metagene MNase-seq analysis of (A) stable and (B) fragile nucleosomes upon RSC and TBP depletion, averaged over all genes as annotated by (Cherry et al., 2012), aligned at the location of the +1 nucleosome (black dashed line).

(B),(E) Heatmap of the log2-fold-change in nucleosome coverage of (B) stable and (E) fragile nucleosomes at all genes as annotated by (Cherry et al., 2012), sorted by NDR width and aligned at the location of the +1 nucleosome, upon RSC and TBP depletion.

(C),(F) Overlay of metagene MNase-seq profiles of (C) stable and (F) fragile nucleosomes upon RSC depletion and simultaneous RSC and TBP depletion, averaged over all genes as annotated by (Cherry et al., 2012), aligned at the location of the +1 nucleosome (black dashed line) .

(G) Cumulative distribution of coverage of stable nucleosomes in TATA or TATA-mismatch elements genome-wide upon depletion of RSC, showing a larger increase in coverage at TATA-mismatch elements than at a TATA-elements.

(H) Cumulative distribution of coverage of stable nucleosomes in TATA or TATA-mismatch elements genome-wide upon simultaneous depletion of RSC and TBP.

(I) Overlay of cumulative distributions of coverage of stable nucleosomes in TATA or TATA-mismatch regions genome-wide upon depletion of RSC or simultaneous depletion of RSC and TBP, showing that partial TBP depletion specifically increased the coverage at TATA-elements.

Shown is one representative replicate out of two experiments for all plots.

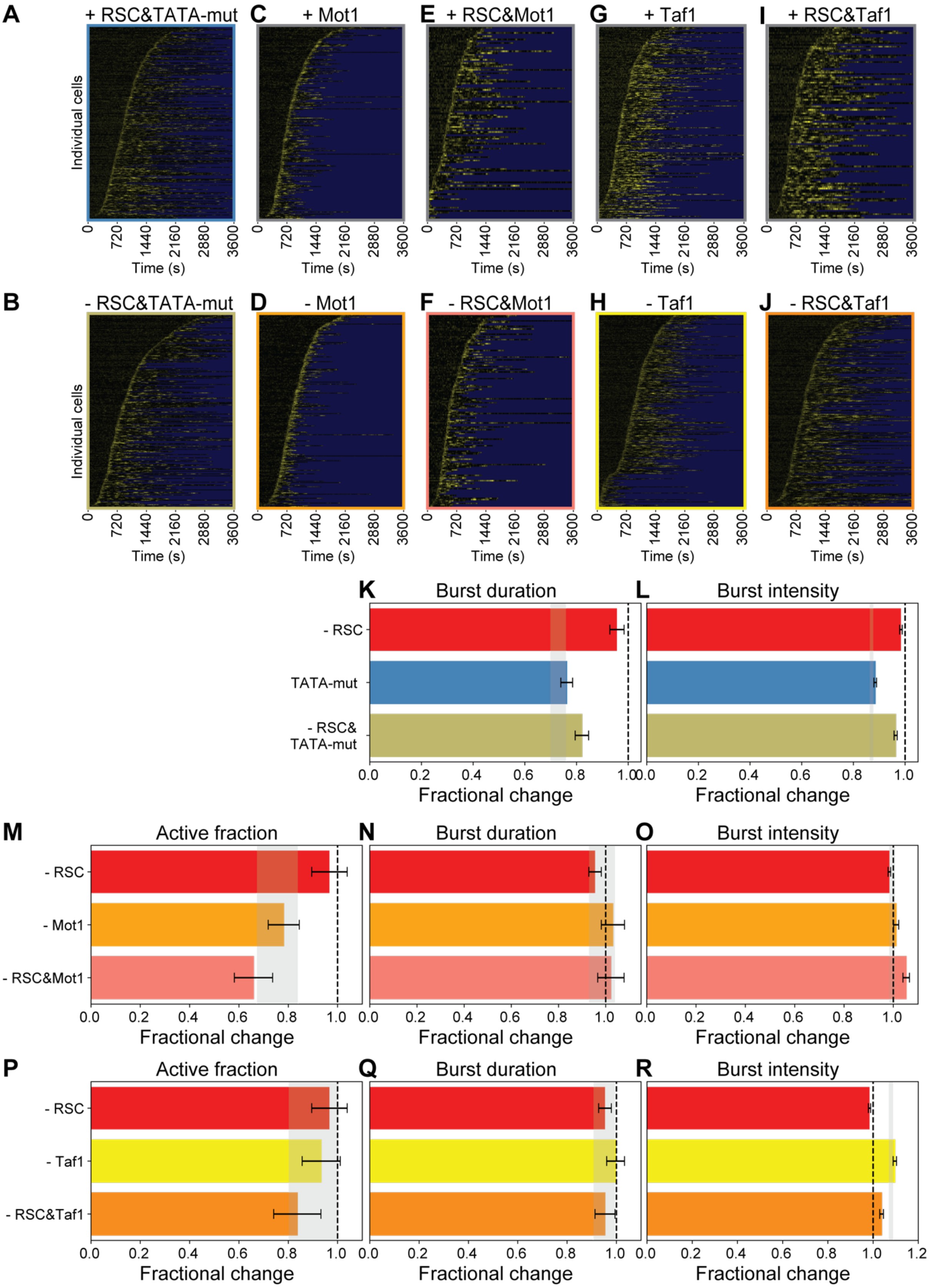


**Figure S7, related to Figure 5 and Figure 6**

**Change in transcriptional bursting when TBP residence time is perturbed in combination with RSC depletion.**

(A-J) Heatmap of the TS intensity in (A) N=315, (B) N=303, (D) N=276, (D) N=285, (E) N=116, (F) N=145, (G) N=231, (H) N=352, (I) N=114 and (J) N=275 cells (rows) in the (A) presence and (B) absence of indicated factors. Yellow: fluorescence intensity of TS; blue: region excluded from analysis.

(K,L) Change in (K) burst duration (and (L) burst intensity when depleting RSC and/or mutating the TATA element. Grey bar: expected effect based on dynamic epistasis analysis.

(M-O) Change in (M) the fraction of cells that activate during 1 hour of imaging, (N) the burst duration and (O) the burst intensity when depleting RSC and/or Mot1. Grey bar: expected effect based on dynamic epistasis analysis.

(P-R) Change in (P) the fraction of cells that activate during 1 hour of imaging, (Q) the burst duration and (R) the burst intensity when depleting RSC and/or Taf1. Grey bar: expected effect based on dynamic epistasis analysis.

Error bars are propagated standard deviations based on 1000 bootstrap repeats.

**Table S1: Yeast strains used in this study**

| YEAST STRAIN | SOURCE |
|---|---|
| S. cerevisiae BY4742: MATα his3Δ1 leu2Δ0 lys2Δ0 ura3Δ0 | Euroscarf |
| S. cerevisiae BY4743: MATa/α his3Δ1/his3Δ1 leu2Δ0/leu2Δ0 LYS2/lys2Δ0 met15Δ0/MET15 ura3Δ0/ura3Δ0 | Euroscarf |
| YTL559: BY4742 gal4::KanMX | this study |
| YTL047: BY4743 with 14xPP7 at 5' GAL10 | Donovan et al, 2019 |
| BY4741 anchor-away (AA) background: tor1-1 fpr1Δ RPL13A-FKBP12-NAT | de Jonge et al, 2017 |
| BY4742 anchor-away (AA) background: tor1-1 fpr1Δ RPL13A-FKBP12-NAT | de Jonge et al, 2017 |
| YTL524: BY4742 AA background with Swi2-FRB-yEGFP1-hphMX4 | kind gift from Holstege lab |
| YTL525: BY4742 AA background with Sth1-FRB-yEGFP1-hphMX4 | kind gift from Holstege lab |
| YTL526: BY4742 AA background with Isw2-FRB-yEGFP1-hphMX4 | kind gift from Holstege lab |
| YTL527: BY4742 AA background with Isw1-FRB-yEGFP1-hphMX4 | kind gift from Holstege lab |
| YTL528: BY4742 AA background with Ino80-FRB-yEGFP1-hphMX4 | kind gift from Holstege lab |
| YTL529: BY4742 AA background with Swr1-FRB-yEGFP1-hphMX4 | kind gift from Holstege lab |
| YTL1178: BY4743 AA background with Sth1-FRB-mScarlet-hphMX4/Sth1-FRB-mScarlet-hphMX4 + GAL10/14xPP7 5'GAL10 + ura3Δ0/ura3-pRPL15A-PCP-GFPEnvy | this study |
| YTL1179: BY4743 AA background with Swi2-FRB-mScarlet-hphMX4/Swi2-FRB-mScarlet-hphMX4 + GAL10/14xPP7 5'GAL10 + ura3Δ0/ura3-pRPL15A-PCP-GFPEnvy | this study |
| YTL1306: BY4742 AA background with Sth1-FRB-yEGFP1-hphMX4 and Swi2-FRB-mScarlet | this study |

| | |
|---|---|
| YTL1309: BY4743 AA background with Sth1-FRB-mScarlet-hphMX4/Sth1-FRB-mScarlet-hphMX4 + Swi2-FRB-mScarlet/Swi2-FRB-mScarlet + GAL10/14xPP7 5'GAL10 + ura3Δ0/ura3-pRPL15A-PCP-GFPEnvy | this study |
| YTL1281: BY4743 AA background with Sth1-FRB-mScarlet-hphMX4/Sth1-FRB-mScarlet-hphMX4 + GAL4/gal4Δ + GAL10/14xPP7 5'GAL10 + ura3Δ0/ura3-pRPL15A-PCP-GFPEnvy | this study |
| YTL1397:BY4742 AA background with Spt15-FRB-mScarlet | this study |
| YTL1505: BY4743 AA background with Spt15-FRB-mScarlet/Spt15-FRB-mScarlet + GAL10/14xPP7 5'GAL10 + ura3Δ0/pRPL15A-PCP-GFPEnvy | this study |
| YTL1506: BY4743 AA background with SPT15/Spt15-FRB-mScarlet + GAL10/14xPP7 5'GAL10 + ura3Δ0/ura3-pRPL15A-PCP-GFPEnvy | this study |
| YTL1584: BY4743 AA background with Sth1-FRB-mScarlet-hphMX4/Sth1-yEGFP1-hphMX4 + Spt15/Spt15-FRB-mScarlet + ura3Δ0/pRPL15A-PCP-GFPEnvy | this study |
| YTL1508: BY4743 AA background with Sth1-FRB-mScarlet-hphMX4/Sth1-FRB-mScarlet-hphMX4 + SPT15/Spt15-FRB-mScarlet + GAL10/14xPP7 5'GAL10 + ura3Δ0/ura3-pRPL15A-PCP-GFPEnvy | this study |
| YTL1510: BY4743 AA background with Swi2-FRB-mScarlet-hphMX4/Swi2-FRB-mScarlet-hphMX4 + with SPT15/Spt15-FRB-mScarlet + GAL10/14xPP7 5'GAL10 ura3Δ0/ura3-pRPL15A-PCP-GFPEnvy | this study |
| YTL1613: BY4742 AA background with TATA-mut-int2-GAL10 | this study |
| YTL1615: BY4742 AA background with Sth1-FRB-yEGFP1-hphMX4 + TATA-mut-int2 | this study |
| YTL1626: BY4743 AA background with Sth1-FRB-mScarlet-hphMX4/Sth1-FRB-mScarlet-hphMX4 + GAL10/TATA-mut-int2 14xPP7 5'GAL10 + ura3Δ0/ura3-pRPL15A-PCP-GFPEnvy | this study |
| YTL1394: BY4742 AA background with Mot1-FRB-mScarlet | this study |
| YTL1470: BY4743 AA background with Mot1-FRB-mScarlet/Mot1-FRB-mScarlet + GAL10/14xPP7 5'GAL10 + ura3Δ0/pRPL15A-PCP-GFPEnvy | this study |
| YTL1588: BY4742 AA background with Sth1-FRB-yEGFP1-hphMX4 + Mot1-FRB-mScarlet | this study |
| YTL1591: BY4743 AA background with Sth1-FRB-mScarlet-hphMX4/Sth1-FRB-mScarlet-hphMX4 + Mot1-FRB-mScarlet/Mot1-FRB-mScarlet + GAL10/14xPP7 5'GAL10 + ura3Δ0/ura3-pRPL15A-PCP-GFPEnvy | this study |
| YTL1391: BY4742 AA background with Taf1-FRB-mScarlet | this study |

| | |
|---|---|
| YTL1448: BY4743 AA background with Taf1-FRB-mScarlet/Taf1-FRB-mScarlet with GAL10/14xPP7 5'GAL10 + ura3Δ0/pRPL15A-PCP-GFPEnvy | this study |
| YTL1413: BY4742 AA background with Sth1-FRB-yEGFP1-hphMX4 + Taf1-FRB-mScarlet | this study |
| YTL1450: BY4743 AA background with Sth1-FRB-mScarlet-hphMX4/Sth1-FRB-mScarlet-hphMX4 + Taf1-FRB-mScarlet/Taf1-FRB-mScarlet + GAL10/14xPP7 5'GAL10 + ura3Δ0/ura3-pRPL15A-PCP-GFPEnvy | this study |

**Table S2: Plasmids used in this study**

| PLASMID | SOURCE |
|---|---|
| pTL100: pFA6a-FRB-yEGFP1-hphMX4 | kind gift from Holstege lab |
| pTL329: pFA6a-FRB-mScarlet-hphMX4 | this study |
| pTL031: 14x PP7 with loxP-kanMX-loxP | Donovan et al 2019 |
| pTL014: pURA PGAL CRE recombinase | Lenstra et al, 2015 |
| pTL191: pHIS PGAL CRE recombinase | this study |
| pTL174: pURA SIV pRPL15A PCP-NLS-GFPEnvy | this study |
| pTL131: pML104 Cas9 guide RNA construct | Laughery et al., 2015 |
| pTL416: pML104 with guideRNA to 3'end SWI2 (gtacttatattgctttagga) | this study |
| pTL363: pML104 with guideRNA to GAL4 (ggtgaaggccctactgagcc) | this study |
| pTL459: pML104 with guideRNA to 3'end SPT15 (gagtagacgaaaagaaaaaa) | this study |
| pTL440: pML104 with guideRNA to 3'end MOT1 (cgataaacaaattcttgcat) | this study |
| PTL126: pML104 with guideRNA to GAL10 TATA element (catataagtaagattagata) | this study |
| pTL444: pML104 with guideRNA to 3'end TAF1 (atcgaataaaacttttgtaa) | this study |

**Table S3: Oligos used in this study**

| OLIGO | SOURCE |
|---|---|
| Sth1-FRB-mScarlet-hphMX4-tag F: tacgttgatgctgacaagttaaatgagtttactgatgaatggttcaaggaacactcttcgcggatccccgggttaattaa | IDT |
| Sth1-FRB-mScarlet-hphMX4-tag R: tattagaggggaaagggatatagtcgtaaaaaaaaaaaacatgtggtgatgaaaacgttagaattcgagctcgtttaaac | IDT |
| Swi2-FRB-mScarlet-hphMX4-tag F or Swi2-FRB-mScarlet-tag F: tcaagcgtggctgaatctttcacagatgaagcggactcgagcatgacagaagcgagtgtacggatccccgggttaattaa | IDT |

| | |
|---|---|
| Swi2-FRB-mScarlet-hphMX4-tag R:ggattaatgtttgtctacgtataaacgaataagtacttatattgctttaggaaggtactagaattcgagctcgtttaaac | IDT |
| Swi2-FRB-mScarlet-tag R: aaaaagagggattaatgtttgtctacgtataaacgaataagtacttatattgctttaggagctaaatgtacgggcgacag | IDT |
| Taf1-FRB-mScarlet-tag F: attagaacaaataaatcttgtccaatgtatagcagtaaagataaccctgcttcaccaaagcggatccccgggttaattaa | IDT |
| Taf1-FRB-mScarlet-tag R: tgatcactgccagtgtttagaaacatagtgtacatttctttattgttacattatacaaaagctaaatgtacgggcgacag | IDT |
| Spt15-FRB-mScarlet-tag F: gaagaaatttaccaagcttttgaagctatataccctgtgctaagtgaatttagaaaaatgcggatccccgggttaattaa | IDT |
| Spt15-FRB-mScarlet-tag R: ccattaataattgagaaaatggaacaaatagaaaaccttttttttcttttcgtctactcctgctaaatgtacgggcgacag | IDT |
| Mot1-FRB-mScarlet-tag F: tgggacccatctcaatacgaggaggagtataatttagacaccttcatcaaaactttacgacggatccccgggttaattaa | IDT |
| Mot1-FRB-mScarlet-tag R: gttgataatacataaatgcgttaaataaaacaaaaatgaccttgtatacgcgtcattccagctaaatgtacgggcgacag | IDT |
| gal4del repair: acgccatcattttaagagaggacagagaagcaagcctcctgaaagaatgaatcgtagatactgaaaaaccccgcaagttcacttcaactg | IDT |
| gal4-deletion-F: tgggacttgaacagctcctt | IDT |
| gal4-deletion-R: tttgggtgtcttcatcacca | IDT |
| TATA-mut-int2 repair: gataatggggctctttacatttccacaagctataagtaagattagatatggatatgtatatggtggtaatgccatgtaatatgattatta | IDT |
| Primer 1.1 Illumina: aatgatacggcgaccaccgagat | IDT |
| Primer 2.1 Illumina: caagcagaagacggcatacga | IDT |

**Table S4: smFISH probes used in this study**

| PROBE | SOURCE |
|---|---|
| PP7-Cy3 smFISH probe 1: [Cy3]atatcgtctgctcctttcta | IDT |
| PP7-Cy3 smFISH probe 2: [Cy3]atatgctctgctggtttcta | IDT |
| PP7-Cy3 smFISH probe 3: [Cy3]gcaattaggtaccttaggat | IDT |
| PP7-Cy3 smFISH probe 4: [Cy3]aatgaacccgggaatactgc | IDT |
| PP7-Cy5 smFISH probe 1: [Cy5]atatcgtctgctcctttcta | IDT |
| PP7-Cy5 smFISH probe 2: [Cy5]atatgctctgctggtttcta | IDT |
| PP7-Cy5 smFISH probe 3: [Cy5]gcaattaggtaccttaggat | IDT |
| PP7-Cy5 smFISH probe 4: [Cy5]aatgaacccgggaatactgc | IDT |

| | |
|---|---|
| GAL10-yeast-1: [Quasar670]aggagtcttcaacctgcaa | Biosearch Technologies |
| GAL10-yeast-2: [Quasar670]aaaggattctcagtagtcca | Biosearch Technologies |
| GAL10-yeast-3: [Quasar670]tggcctcgacaccccttaa | Biosearch Technologies |
| GAL10-yeast-4: [Quasar670]catatcttcagcggaaaatc | Biosearch Technologies |
| GAL10-yeast-5: [Quasar670]atagtcacaaatcttgcgtc | Biosearch Technologies |
| GAL10-yeast-6: [Quasar670]ggcttgaaatctggtgccg | Biosearch Technologies |
| GAL10-yeast-7: [Quasar670]tcactttcaggtcaacaatg | Biosearch Technologies |
| GAL10-yeast-8: [Quasar670]atagccaagaacaactgatt | Biosearch Technologies |
| GAL10-yeast-9: [Quasar670]gaattcgacaggttatcagc | Biosearch Technologies |
| GAL10-yeast-10: [Quasar670]caaatacccttcctcatttt | Biosearch Technologies |
| GAL10-yeast-11: [Quasar670]gcgcctatataagcactatc | Biosearch Technologies |
| GAL10-yeast-12: [Quasar670]catacctgccgatcgtg | Biosearch Technologies |
| GAL10-yeast-13: [Quasar670]actaaacttacccttcgaaa | Biosearch Technologies |
| GAL10-yeast-14: [Quasar670]acggttaactgatagtcttt | Biosearch Technologies |
| GAL10-yeast-15: [Quasar670]ctatgattcgcattaacgcc | Biosearch Technologies |
| GAL10-yeast-16: [Quasar670]tctgtggaaagaaccgatac | Biosearch Technologies |
| GAL10-yeast-17: [Quasar670]aaggattttgaatgatgggt | Biosearch Technologies |
| GAL10-yeast-18: [Quasar670]cctggctacagaatcataag | Biosearch Technologies |
| GAL10-yeast-19: [Quasar670]atgtactcggcggtaaaa | Biosearch Technologies |
| GAL10-yeast-20: [Quasar670]tcggtgtccttctcattatc | Biosearch Technologies |
| GAL10-yeast-21: [Quasar670]ttaccaatagatcacctgga | Biosearch Technologies |
| GAL10-yeast-22: [Quasar670]ttgggcaacgttcacagtat | Biosearch Technologies |
| GAL10-yeast-23: [Quasar670]ccagcagtcaatttaccttt | Biosearch Technologies |
| GAL10-yeast-24: [Quasar670]ttaaatttattggcgtcgct | Biosearch Technologies |
| GAL10-yeast-25: [Quasar670]cctcaatagtgtctccatat | Biosearch Technologies |
| GAL10-yeast-26: [Quasar670]ttgaacgcaccataatctcc | Biosearch Technologies |
| GAL10-yeast-27: [Quasar670]attacccgtaggaatcatgt | Biosearch Technologies |
| GAL10-yeast-28: [Quasar670]ggctttgtagagttaaaggt | Biosearch Technologies |
| GAL10-yeast-29: [Quasar670]acaatcaaactggggatttt | Biosearch Technologies |
| GAL10-yeast-30: [Quasar670]ttgacttggcttagcatttt | Biosearch Technologies |
| GAL10-yeast-31: [Quasar670]aacctcatagaagggaatgt | Biosearch Technologies |
| GAL10-yeast-32: [Quasar670]atcgggatgaaaagccttga | Biosearch Technologies |
| GAL10-yeast-33: [Quasar670]tggctctgtacttaaaact | Biosearch Technologies |
| GAL10-yeast-34: [Quasar670]cagcagacaagaaatcaccg | Biosearch Technologies |
| GAL10-yeast-35: [Quasar670]accttgtcttgcttcgtaac | Biosearch Technologies |
| GAL10-yeast-36: [Quasar670]aatgtatctaccaggctcaa | Biosearch Technologies |
| GAL10-yeast-37: [Quasar670]ctttgtaactgagctgtcat | Biosearch Technologies |
| GAL10-yeast-38: [Quasar670]acacaatctttccagttctc | Biosearch Technologies |
| GAL10-yeast-39: [Quasar670]ccttttcggtcacacaaatc | Biosearch Technologies |
| GAL10-yeast-40: [Quasar670]caatcttggacccgtaagtt | Biosearch Technologies |
| GAL10-yeast-41: [Quasar670]agtgaattaccgaatcaatttta | Biosearch Technologies |
| GAL10-yeast-42: [Quasar670]cacctacagcctttaaacca | Biosearch Technologies |

| | |
|---|---|
| GAL10-yeast-43: [Quasar670]gtgatagtatctcagcggga | Biosearch Technologies |
| GAL10-yeast-44: [Quasar670]cctgtaaccaaaacaattttaga | Biosearch Technologies |
| GAL10-yeast-45: [Quasar670]ctaataaaacgacagttccc | Biosearch Technologies |
| GAL10-yeast-46: [Quasar670]atttggaaacgttgtattgt | Biosearch Technologies |
| GAL10-yeast-47: [Quasar670]caccatagacagtagcagaa | Biosearch Technologies |
| GAL10-yeast-48: [Quasar670]atttgggaatctcgtagcat | Biosearch Technologies |

**Table S5: Sequencing adapters used in this study**

| SEQUENCING ADAPTER | SOURCE |
|---|---|
| Universal adapter: aatgatacggcgaccaccgagatctacactctttccctacacgacgctcttccgatc*t | IDT |
| Sequencing adapter 001: 5'-Phos-gatcggaagagcacacgtctgaactccagtcacatcacgatctcgtatgccgtcttctgcttg | IDT |
| Sequencing adapter 002: 5'-Phos-gatcggaagagcacacgtctgaactccagtcaccgatgtatctcgtatgccgtcttctgcttg | IDT |
| Sequencing adapter 003: 5'-Phos-gatcggaagagcacacgtctgaactccagtcacttaggcatctcgtatgccgtcttctgcttg | IDT |
| Sequencing adapter 004: 5'-Phos-gatcggaagagcacacgtctgaactccagtcactgaccaatctcgtatgccgtcttctgcttg | IDT |
| Sequencing adapter 005: 5'-Phos-gatcggaagagcacacgtctgaactccagtcacacagtgatctcgtatgccgtcttctgcttg | IDT |
| Sequencing adapter 006: 5'-Phos-gatcggaagagcacacgtctgaactccagtcacgccaatatctcgtatgccgtcttctgcttg | IDT |
| Sequencing adapter 007: 5'-Phos-gatcggaagagcacacgtctgaactccagtcaccagatcatctcgtatgccgtcttctgcttg | IDT |
| Sequencing adapter 008: 5'-Phos-gatcggaagagcacacgtctgaactccagtcacacttgaatctcgtatgccgtcttctgcttg | IDT |
| Sequencing adapter 009: 5'-Phos-gatcggaagagcacacgtctgaactccagtcacgatcagatctcgtatgccgtcttctgcttg | IDT |
| Sequencing adapter 010: 5'-Phos-gatcggaagagcacacgtctgaactccagtcactagcttatctcgtatgccgtcttctgcttg | IDT |
| Sequencing adapter 012: 5'-Phos-gatcggaagagcacacgtctgaactccagtcaccttgtaatctcgtatgccgtcttctgcttg | IDT |
| Sequencing adapter 013: 5'-Phos-gatcggaagagcacacgtctgaactccagtcacagtcaaatctcgtatgccgtcttctgcttg | IDT |
| Sequencing adapter 014: 5'-Phos-gatcggaagagcacacgtctgaactccagtcacagttccatctcgtatgccgtcttctgcttg | IDT |
| Sequencing adapter 015: 5'-Phos-gatcggaagagcacacgtctgaactccagtcacatgtcaatctcgtatgccgtcttctgcttg | IDT |
| Sequencing adapter 016: 5'-Phos-gatcggaagagcacacgtctgaactccagtcacccgtccatctcgtatgccgtcttctgcttg | IDT |

Video S1
Related to Figure 1
Fluorescence imaging of a TS showing *GAL10* transcription dynamics in a representative cell in the presence of RSC. Cells were imaged every 15 s for 1 hour after galactose addition.

Green box: location of *GAL10* TS. Shown is a maximum intensity projection (top left) with *xz* (bottom) and *yz* (right) sideviews.
